# Bacterial analogs of plant piperidine alkaloids mediate microbial interactions in a rhizosphere model system

**DOI:** 10.1101/499731

**Authors:** Gabriel L. Lozano, Hyun Bong Park, Juan I. Bravo, Eric A. Armstrong, John M. Denu, Eric V. Stabb, Nichole A. Broderick, Jason M. Crawford, Jo Handelsman

**Author notes:** Co-correspondence.

## Abstract

Plants expend significant resources to select and maintain rhizosphere communities that benefit their growth and protect them from pathogens. A better understanding of assembly and function of rhizosphere microbial communities will provide new avenues for improving crop production. Secretion of antibiotics is one means by which bacteria interact with neighboring microbes and sometimes change community composition. In our analysis of a taxonomically diverse consortium from the soybean rhizosphere, we found that *Pseudomonas koreensis* selectively inhibits growth of *Flavobacterium johnsoniae* and other members of the Bacteroidetes grown in soybean root exudate. A genetic screen in *P. koreensis* identified a previously uncharacterized biosynthetic gene cluster responsible for the inhibitory activity. The metabolites were isolated based on biological activity and were characterized using tandem-mass spectrometry, multidimensional NMR, and Mosher ester analysis, leading to the discovery of a new family of bacterial piperidine alkaloids, koreenceine A-D (**1–4**). Three of these metabolites are analogs of the plant alkaloid γ-coniceine. Comparative analysis of the koreenceine cluster with the γ-coniceine pathway revealed distinct polyketide synthase (PKS) routes to the defining piperidine scaffold, suggesting convergent evolution. Koreenceine-type pathways are widely distributed among *Pseudomonas* species, and koreenceine C was detected in another *Pseudomonas* sp. from a distantly related cluster. This work suggests that *Pseudomonas* and plants convergently evolved the ability to produce similar alkaloid metabolites that can mediate inter-bacterial competition in the rhizosphere.

**IMPORTANCE:** The microbiomes of plants are critical to host physiology and development. Microbes are attracted to the rhizosphere due to massive secretion of plant photosynthates from roots. Microorganisms that successfully join the rhizosphere community from bulk soil have access to more abundant and diverse molecules, producing a highly competitive and selective environment. In the rhizosphere, as in other microbiomes, there is little known about the genetic basis for individual species’ behaviors within the community. In this study, we characterized competition between *Pseudomonas koreensis* and *Flavobacterium johnsoniae*, two common rhizosphere inhabitants. We identified a widespread gene cluster in several *Pseudomonas* spp., which is necessary for the production of a novel family of piperidine alkaloids that are structural analogs of plant alkaloids. We expand the known repertoire of antibiotics produced from *Pseudomonas* in the rhizosphere and demonstrate the role of the metabolites in interactions with other bacteria of the rhizosphere.

## INTRODUCTION

Plants were long thought to be defined by their genes and environments. It has recently become apparent that plants are also shaped by their microbiomes — the communities of microorganisms that live on, around, and inside them (1). Microbiomes modify many environments, including humans, animals, oceans, soils, and hot springs. Comprehensive investigations of the interactions between microbiomes and their environments, as well as the interactions within microbiomes that contribute to their function and stability, are important to understand diverse niches on Earth, including those associated with plants.

The rhizosphere comprises plant root surfaces and their surrounding soil microenvironments. Bacteria are attracted to this environments by the massive amount of plant photosynthate, in the form of sugars, organic acids, and amino acids, which is secreted from roots (2). Bacteria that colonize the rhizosphere play an essential role in plant growth, and resistance to pathogens. For example, some members secrete plant-like hormones, such as indole acetic acid, gibberellic acid, cytokinin and abscisic acid, that promote plant growth (3), whereas others suppress plant diseases by secreting diverse compounds such as zwittermicin A, 2,4-diacetylphloroglucinol, and pyoluteorin (4). Thus, bacterial rhizosphere communities represent a rich reservoir of bioactive metabolites.

Use of bacteria for biological control of plant disease has been pursued for decades, but foreign microorganisms typically do not persist in native rhizosphere communities (5). Nutrient abundance, host availability, and microbial interactions define indigenous microbial community structures and limit colonization by invading bacteria. To engineer plant microbiomes to improve agricultural systems, a better understanding of the inter-bacterial interactions that dominate the rhizosphere is needed.

We developed ***t***he ***h***itchhikers ***o***f the ***r***hizosphere (THOR), a model system to examine the molecular interactions among core bacterial members of the rhizosphere (6). This model system is composed of *Bacillus cereus, Flavobacterium johnsoniae*, and *Pseudomonas koreensis*, which belong to three dominant phyla within the rhizosphere — Firmicutes, Bacteroidetes, and Proteobacteria, respectively. The three members display both competitive and cooperative interactions. For example, *P. koreensis* inhibits growth of *F. johnsoniae*, but not in the presence of *B. cereus*. Inhibition was only observed when bacteria were grown in soybean root exudate and is specific for Bacteroidetes, since this phylum was the only one inhibited by *P. koreensis* from a collection of taxonomically diverse rhizosphere bacteria (6). In this study, we characterized the genetic and molecular mechanisms by which *P. koreensis* inhibits *F. johnsoniae*. We determined that a new family of bacterial piperidine alkaloids designated koreenceine A-D (**1–4**) are produced by an orphan polyketide synthase (PKS) pathway and mediate inhibition of members of the Bacteroidetes. Koreenceines A, B and C are structural analogs of the piperidine alkaloid γ-coniceine, produced by plants, and comparisons of the plant and bacterial biosynthetic pathways support a convergent evolutionary model.

## RESULTS

### Identification of an orphan *P. koreensis* pathway that is responsible for inhibiting growth of *F. johnsoniae*

To identify the genes required for inhibition of *F. johnsoniae* by *P. koreensis* in root exudate, we screened 2,500 *P. koreensis* transposon mutants and identified sixteen that did not inhibit *F. johnsoniae* (Table 1). Two of these mutants mapped to an uncharacterized polyketide biosynthetic cluster containing 11 genes (Fig. 1). We deleted the entire gene cluster (Δ*kecA-kec*F::*tet*), which abolished inhibitory activity against *F. johnsoniae* and other members of the Bacteroidetes (Fig. S1). We designed this pathway as an orphan pathway since the encoded natural product is unknown.

**Table 1.** *P. koreensis* mutants identified in the genetic screen with loss of inhibitory activity against *F. johnsoniae*.

| Mutant | ID | Predicted Function | Class |
| --- | --- | --- | --- |
| 1 | OOH83074 | Pyridoxalphosphate dependent aminotransferase | Secondary metabolites production |
| 2 | OOH83072 | 3-oxoacyl-ACP synthase | Secondary metabolites production |
| 3, 4 | OOH76777 | Two-component sensor histidine kinase CbrA | Cell signaling and transcription regulation |
| 5, 6 | OOH78126 | CysB family transcriptional regulator | Cell signaling and transcription regulation |
| 7 | OOH78405 | Multi-sensor hybrid histidine kinase | Cell signaling and transcription regulation |
| 8 | OOH81884 | Outer membrane protein assembly factor BamC | Cell surface |
| 9 | OOH77437 | Peptidoglycan-associated lipoprotein | Cell surface |
| 10 | OOH75718 | Phospholipid/glycerol acyltransferase | Cell surface |
| 11 | OOH75539 | Acetylglutamate kinase | Metabolism |
| 12 | OOH81129 | Methylcitrate synthase | Metabolism |
| 13 | OOH76758 | Ketol-acid reductoisomerase | Metabolism |
| 14 | OOH80977 | Succinyl-CoA synthetase subunit alpha | Metabolism |
| 15 | OOH81130 | 2-methylisocitrate dehydratase | Metabolism |
| 16<sup>a</sup> | OOH76781 | 3-methyl-2-oxobutanoate hydroxymethyltransferase | Metabolism |
|  | OOH76782 | Pantoate-beta-alanine ligase | Metabolism |

<sup>a</sup> Transposon insertion in the promoter, 5' region, of an operon conformed for these two genes

**FIG 1.**
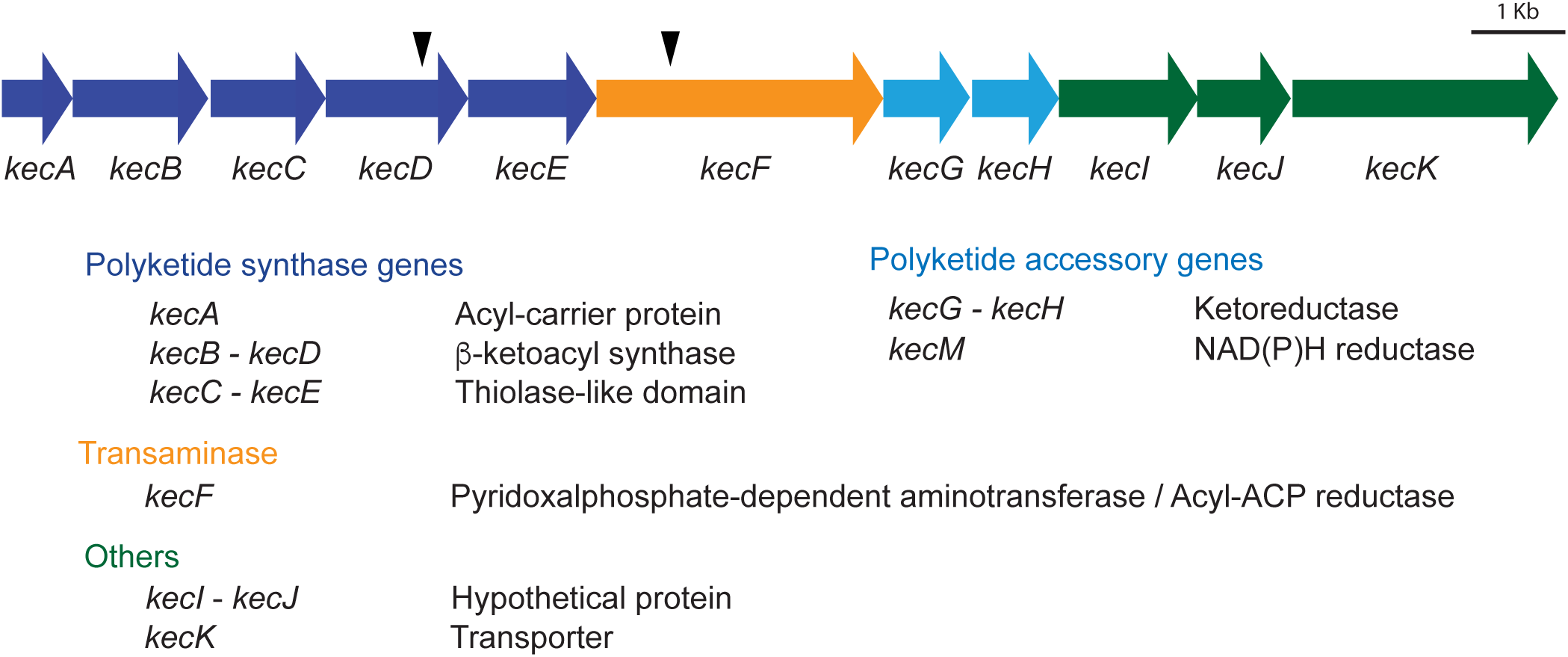
Koreenceine biosynthetic locus and the predicted function of each gene. Black arrows indicate locations of the transposons of the mutants identified.

We developed a defined medium in which *P. koreensis* simulated the gene cluster-dependent inhibitory activity against *F. johnsoniae* that was observed in root exudate (Fig. S2A). Since we identified two independent mutants in the gene encoding a sensor histidine kinase, *cbrA*, that was required for activity in root exudate (Table 1), we developed a defined medium with the goal of activating the CbrAB system (7), which controls the utilization of alternative carbon sources such as amino acids (8). Adding to defined medium the same mix of the amino acids that was used to supplement root exudate induced *P. koreensis* to produce inhibitory activity in the defined media (Fig. S2A). We tested 19 different individual amino acids, and identified five, including aspartate, that induce inhibition of *F. johnsoniae* by *P. koreensis* (Fig. S2B). A non-hydrolysable analog of aspartate, *N*-methyl-DL-aspartate (Asp*), did not stimulate inhibitory activity, suggesting that catabolism of certain amino acids is required for activity (Fig. S2B).

### Characterization of koreenceine metabolites from the orphan *P. koreensis* pathway

To characterize the inhibitory metabolites from the orphan *P. koreensis* pathway, we compared the metabolomes of the wild-type strain and the non-inhibitory mutant grown in root exudate. High-performance liquid chromatography/mass spectrometry (HPLC/MS)-based analysis of the crude organic extracts led to the identification of peaks **1–4** that were completely abolished in the mutant (Fig. 2). We carried out bioassay-guided preparative-scale HPLC fractionation of the crude organic extract from a culture (5 L) of the wild-type *P. koreensis* grown in defined medium. Peaks **1, 2** and **4** were detected in fractions with antimicrobial activity against *F. johnsoniae*. High-resolution electrospray ionization-quadrupole-time-of-flight MS (HRESIQTOFMS) data of **1–4** revealed *m/z* 208.2067, 210.2224, 226.2171, and 278.1885, allowing us to calculate their molecular formulas as C_14_H_26_N, C_14_H_28_N, C_14_H_27_NO, and C_14_H_29_ClNO_2_, respectively (Fig. 2, Fig. S3). We then proceeded with mass-directed isolation of these metabolites from a larger-scale culture in defined medium (12 L) of wild-type *P. koreensis* for NMR-based structural characterization.

**FIG 2.**
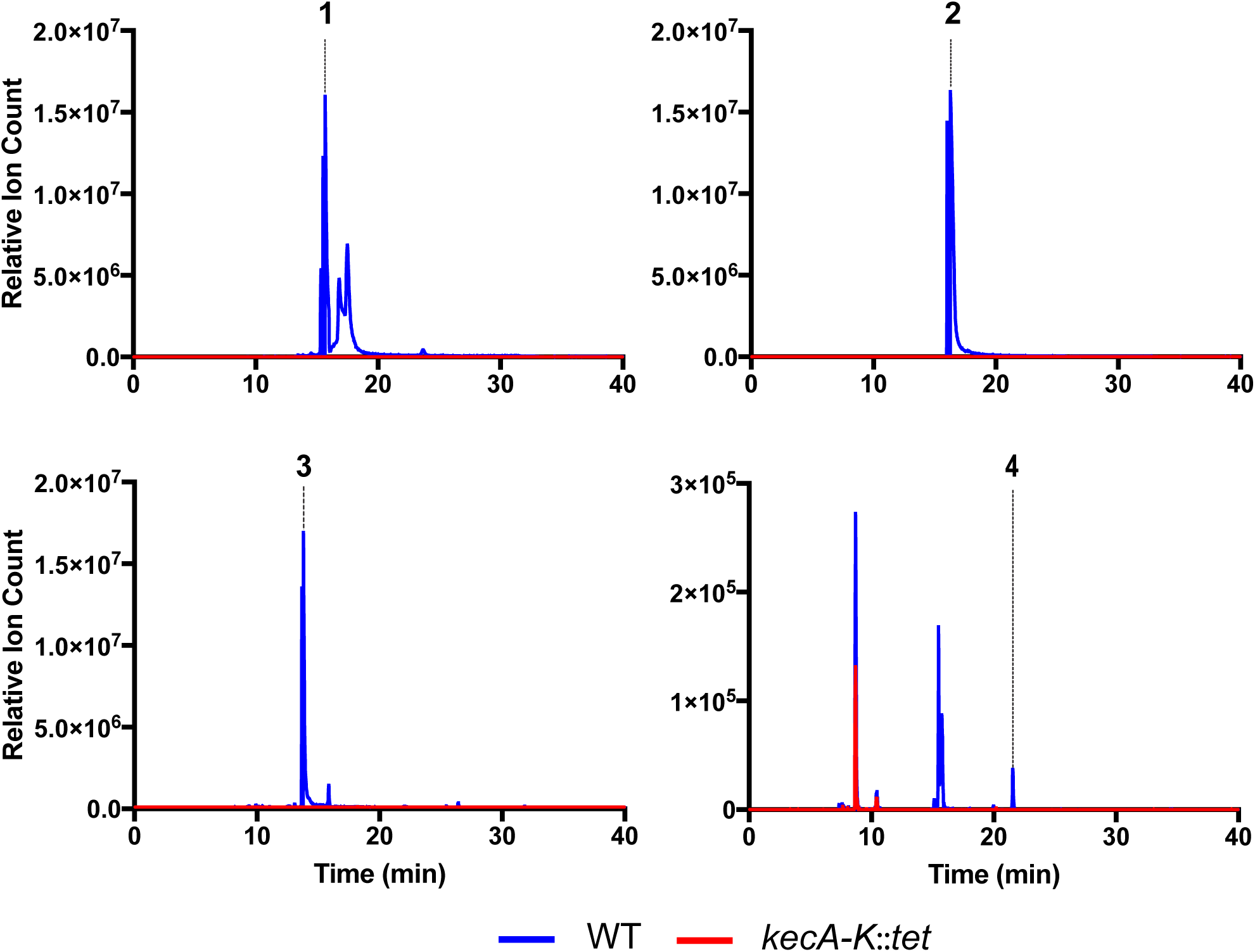
Extracted ion chromatograms of koreenceine A-D for wild type and *kecA-K* deletion mutant.

The chemical structures of **1–4** were characterized through ^1^H, 2D-NMR (gCOSY, gHSQC, and gHMBC), tandem MS, and Mosher ester analysis (Fig. 3, Fig. S3–7). Briefly, ^1^H NMR spectra combined with gHSQC of **2** revealed the presence of six methylene groups including one downfield-shifted signal, six additional methylene, and one methyl group. Consecutive COSY cross-peaks from a triplet methyl H-15 (*δ*_H_ 1.86) to a methylene H-7 (*δ*_H_ 2.53) established a partial structure of a nonane-like hydrocarbon chain. Additional COSY correlations from a downfield-shifted methylene H-2 (*δ*_H_ 3.53) to a methylene H-5 (*δ*_H_ 2.72) also constructed a shorter 4×CH_2_ chain. Key HMBC correlations from H-2, H-4 and H-7 to C-6 allowed us to construct the piperidine core in **2**. In contrast, the ^1^H NMR spectrum of **4** showed the presence of a hydroxyl methine H-3 (*δ*_H_ 3.94), which was evident by COSY correlations with both methylene H-2 and H-4. The connectivity between H-1′ (*δ*_H_ 3.20) and H-4′ (*δ*_H_ 3.59) was established by additional COSY correlations, which was further supported to be a 4-chlorobutanamine-like partial structure by the presence of a mono-chlorine isotope distribution pattern in the HRESIQTOFMS data. HMBC correlations from H-2 and H-1′ to an amide carbon C-1 unambiguously constructed the chemical structure of **4** to be *N*-(4-chlorobutyl)-3-hydroxydecanamide. Modified Mosher’s reaction on the secondary alcohol at C-3 determined the absolute configuration of C-3 to be *R*, completing the absolute structure of 4. The structure of metabolite **1**, an analog of **2**, was elucidated based on the ^1^H and COSY NMR data that indicate the position of a *trans-double* bond between H-7and H-8. Finally, the chemical structure of **3** was deduced by comparative high-resolution tandem MS analyses with the closely related metabolites **1** and **2**.

**FIG 3.**
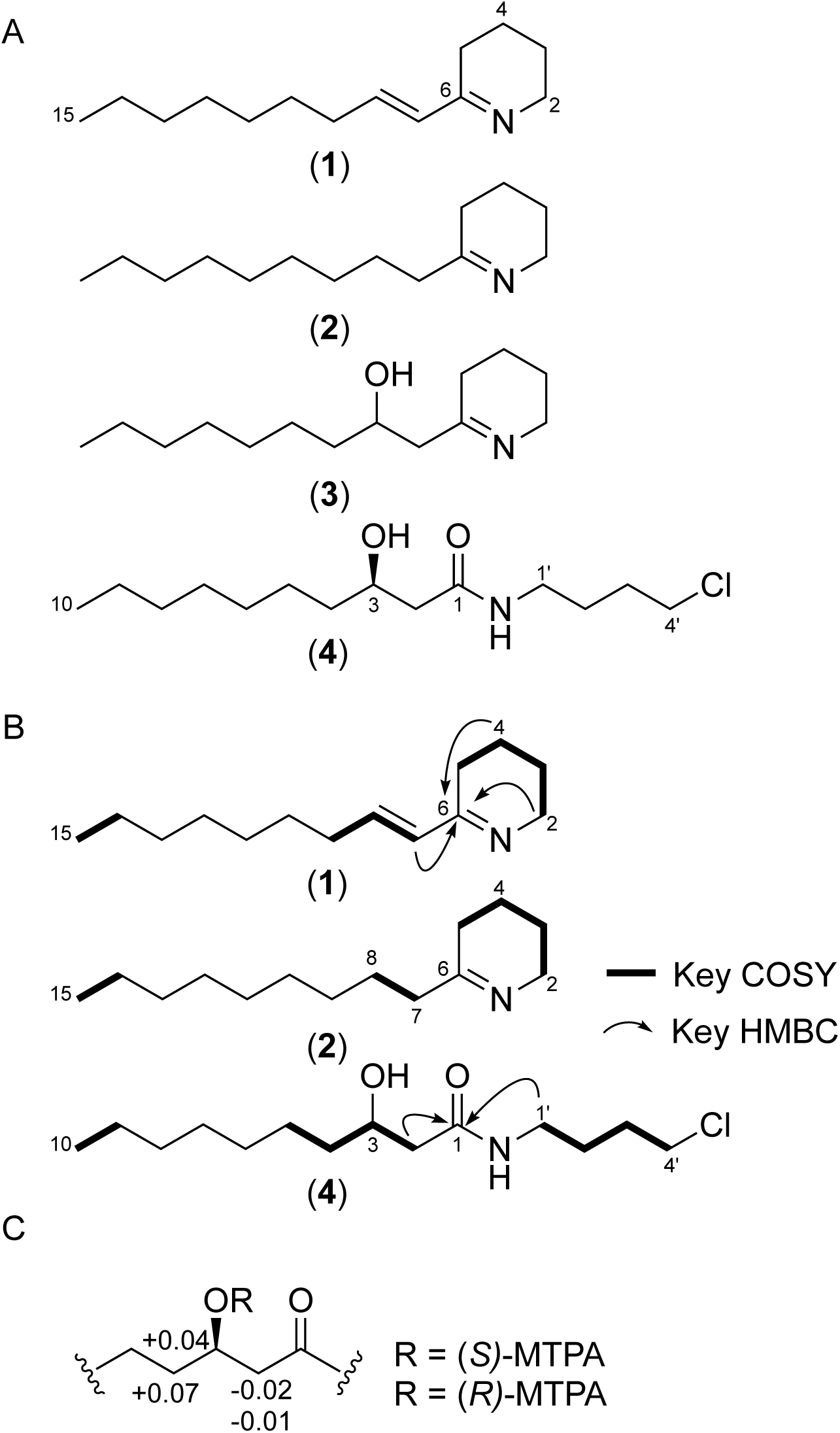
Structural characterization of koreenceines A) Chemical structures of compounds **1–4**. B) Key COSY and HMBC NMR correlations of compounds. C) Δ*δ*_S-R_ in ppm for the MTPA esters of compound **4**.

### Koreenceine structure-activity analysis

We estimated the minimal inhibitory concentration (MIC) values of koreenceine B **(2)** and D **(4)**, as 200 μg mL^-1^ for both metabolites against *F. johnsoniae*. We predicted that koreenceine D does not have a major role in the inhibitory activity, since koreenceine D **(4)** is present in root exudate cultures at levels 100-times less than koreenceine A **(1)**, B **(2)**, and C **(3)** (Fig. 2). We could not estimate an MIC for koreenceine A, as its levels diminish during the purification process. We synthesized koreenceine A and observed similar decomposition during purification (data not shown). Thus, we tested a semi-purified fraction of koreenceine B with a trace of koreenceine A, which had a stronger inhibitory effect than koreenceine B alone, (MIC 40 μg mL^-1^). The significant increase in activity associated with trace amounts of koreenceine A suggests that this molecule is the major inhibitory molecule against *F. johnsionae* in the THOR rhizosphere model or is synergistic with koreenceine B.

### Proposed biosynthesis of koreenceine metabolites

The defining piperidine core of koreenceine metabolites A-C is observed in plant alkaloids such as γ-coniceine, a well characterized alkaloid from poison hemlock *(Conium maculatum)* (9). We analyzed the putative activities of the genes in the koreenceine biosynthetic cluster and identified genes with predicted or previously identified enzymatic activities needed for the production of γ-coniceine in plants (10–13) (Fig. 1, Fig. 4). We propose the following biosynthetic pathway of koreenceine A to C. The first five genes of the cluster, *kecABCDE*, encode a type II polyketide synthase system: *kecA* encodes an acyl carrier protein (ACP); *kecB* and *kecD* encode β-ketoacyl synthases (KSα); and *kecC* and *kecE* encode partial β-ketoacyl synthases with conserved thiolase domains (Chain-Length Factor-CLF or KSβ). This cluster may encode production machinery for two-heterodimer systems, KecB-KecC and KecD-KecE, for polyketide elongation over KecA, and might participate in the formation of a triketide intermediate derived from the condensation of two malonyl units and a decanoyl-, 3-hydroxy-decanoyl-, or a *trans-2*-decenoyl-unit. β-keto reductive modifications could be catalyzed by KecG and KecH reductases (Figure 4). Aminotransferase KecF is predicted to catalyze transamination of the aldehyde intermediate facilitating piperidine cyclization. KecF appears to be a multidomain protein with a predicted aminotransferase at the *N*-terminus and a general NAD(P)-binding domain (IPR036291) and a conserved protein domain COG5322 at the C-terminus. Interestingly, long-chain fatty acyl-ACP reductases from Cyanobacteria share these features and generate fatty aldehydes from the reduction of fatty acid intermediates bound to ACP (14). We predict that KecF reduces the ACP-polyketide intermediate to a polyketide aldehyde with the *C*-terminal domain, followed by transamination by the *N*-terminal domain. Finally, the amine intermediate could undergo a non-enzymatic cyclization as observed in γ-coniceine (12). We predict that koreenceine D is derived from koreenceine C by an unidentified halogenase reaction, as analogous metabolite sets have been detected in plants (Fig. S8) (15). The last three genes, *kecIJK*, may participate in the translocation of the koreenceine alkaloids outside of the cell. KecI and KecJ are hypothetical proteins predicted to localize in the membrane and KecK has homology with membrane-bound drug transporters.

**FIG 4.**
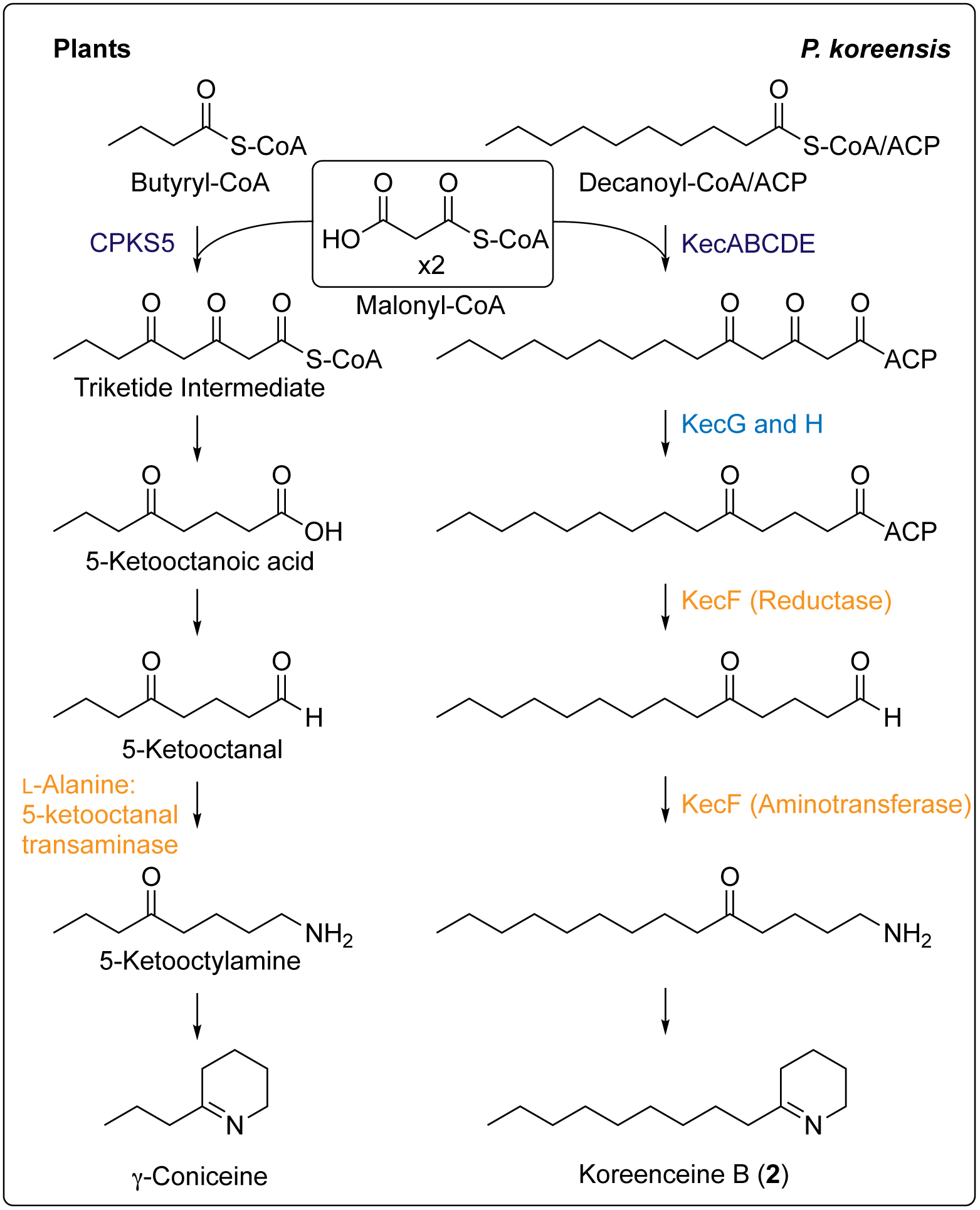
Predicted biosynthetic pathway for γ-coniceine in plants and proposed biosynthetic pathway for koreenceine B in *P. koreensis*. Similar functions are color coded to highlight the similarity between both routes in plants and koreenceine biosynthetic locus.

### Convergent evolution of pathways for production of γ-coniceine-like alkaloids in plants and *P. koreensis*

The biosynthetic pathway for production of γ-coniceine is still under investigation, but ^14^C-feeding experiments in *C. maculatum* coupled with chemical degradation of the labeled products suggest that γ-coniceine is not derived from an amino acid as are other plant alkaloids, but rather it is derived from a polyketide chain produced by the condensation of acetate units (10). Type III polyketide synthases common in plants are iterative homodimers that orchestrate the acyl-CoA mediated priming, extension, and cyclization reactions for polyketide products without the use of acyl carrier proteins (16). Recently, Hotti et al. found CPKS5, a non-chalcone synthase/stilbene synthase (CHS/STS)-type III polyketide synthase expressed in tissues that contain γ-coniceine (Fig. 4) (11). The pathway that we identified in *P. koreensis* predicted two type II PKSs are involved in production of the polyketide intermediate (Fig. 1). Although the pathway that we identified in *P. koreensis* produces compounds related to the plant alkaloids, the PKSs from the plant and bacterial kingdoms share little similarity. We propose that convergent evolution led to two different polyketide pathways for the production of γ-coniceine-like metabolites in plants and bacteria.

### Distribution of the koreenceine cluster

Similar koreenceine-like clusters have previously been identified by functional screens for antimicrobial activities (17, 18); however, there are no reports of the metabolites produced. We identified 179 koreenceine-like clusters in genomes in NCBI (June 2018). The majority of these clusters are in *Pseudomonas* genomes, although we found some partial clusters lacking *kecIJK* in *Xenorhabdus* and *Streptomyces* spp. genomes (Fig. 5A). We used maximum-likelihood analysis of the amino acid sequence of the aminotransferase-reductase protein, KecF, as a representative of the koreenceine cluster for phylogenetic reconstruction. We observed four main clades that are each associated with a bacterial genus (Fig. 5A). Clades A and B, which contain 93% of the clusters, are found in *Pseudomonas* genomes. The koreenceine gene cluster identified in *P. koreensis* in this study belongs to Clade A; other Clade A clusters are located in the same genomic context in *P. koreensis* and *P. mandelii*, two closely related species in the *P. fluorescens* complex (19) (Fig. 5B). Clades C and D were found in *Streptomyces* and *Xenorhabdus* spp., respectively.

**FIG 5.**
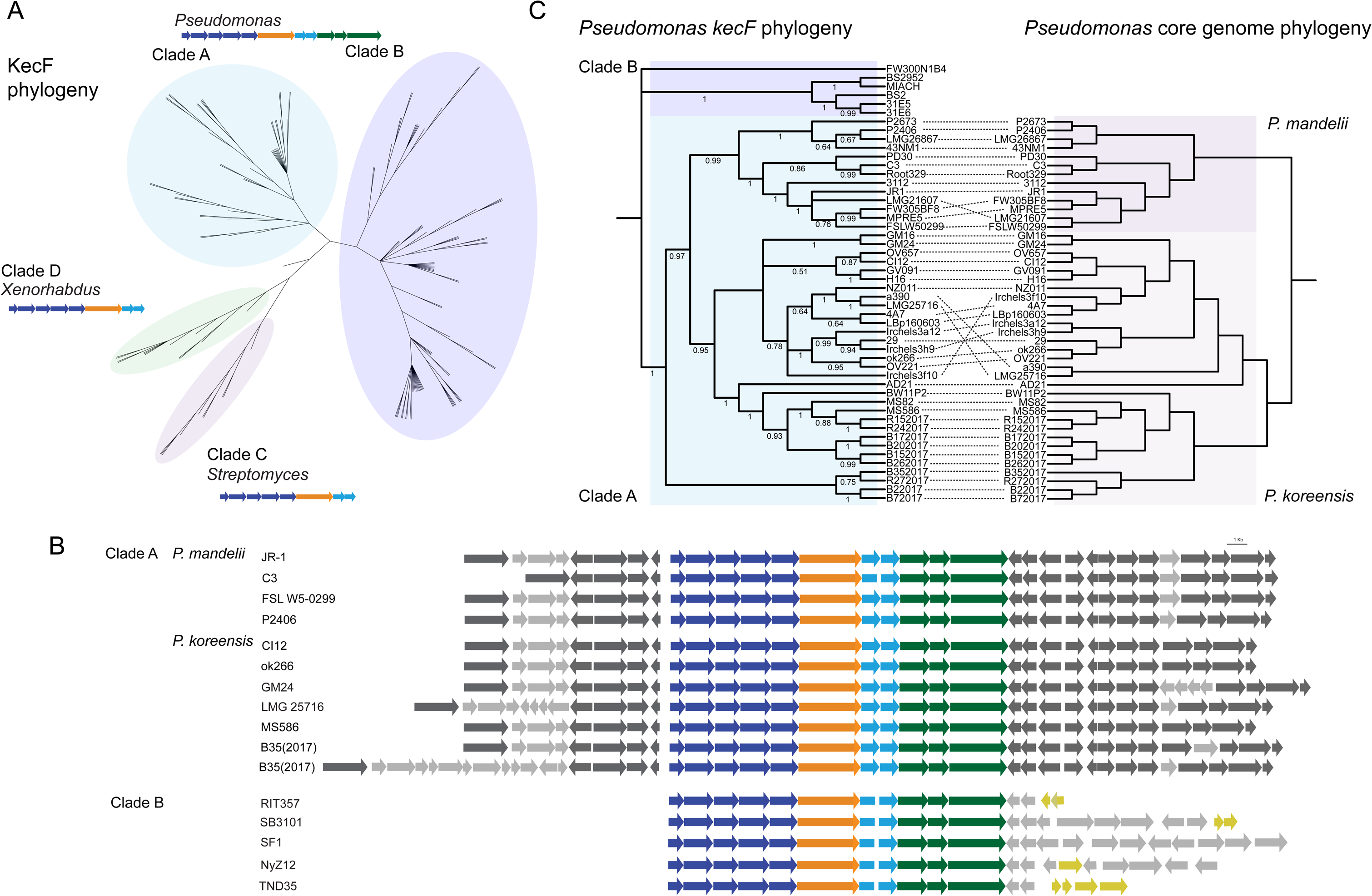
Phylogenetic analysis of the koreenceine biosynthetic locus and its distribution across bacteria. A) Maximum likelihood phylogenetic tree estimated from the amino acid sequence of KecF, and the corresponding structure of the koreenceine-like gene cluster present in each clade. B) Schematic representation of the koreenceine-like biosynthetic locus and its genomic context from several *Pseudomonas* spp. from clade A and clade B. Genes conserved in all genomes from *P. mandelii* and *P. koreensis* are in dark gray, meanwhile variable or unique genes are in gray. Genes that likely experienced horizontal gene transfer events are in yellow. C) Comparison of the phylogenies of *kecF* genes and their associated *Pseudomonas* genomes belonging to Clade A *kecF* homologues. Both phylogenies correspond to maximum likelihood analyses of the nucleotide sequence of the genes or the core genome. The dotted lines connect the cluster with its corresponding *Pseudomonas* genome.

Clade A and the *P. koreensis* and *P. mandelii* genomes are phylogenetically parallel (Fig. 5C). We hypothesize that the gene cluster was acquired before *Pseudomonas fluorescens* complex diversification from a common ancestor of *P. koreensis* and *P. mandelii* and maintained in these *Pseudomonas spp*. by vertical transmission. In contrast, Clade B contains clusters present in *P. putida* and several species of the *P. fluorescens* complex, in which there is no conservation in the genome localization, and the clusters are frequently associated with elements that mediate horizontal gene transfer (Fig. 5B). This suggests different strategies to maintain koreenceine-type gene clusters in diverse *Pseudomonas* species.

Another *Pseudomonas* isolate (SWI36) was reported to inhibit *B. cereus*, and its activity was dependent on a koreenceine-type gene cluster from clade B (18), but we found that it did not inhibit *F. johnsoniae*. Under certain conditions, *P. koreensis* inhibited *B. cereus*, and the activity was dependent on the koreenceine cluster, suggesting a similarity with the SWI36 cluster (Fig. S9). Indeed, targeted metabolomic analysis of *Pseudomonas* sp. SWI36 cell-free culture detected koreenceine C (Fig. S10). These data suggest that different koreenceine-like gene clusters in *Pseudomonas* genomes have the capacity to synthesize koreenceine metabolites.

## DISCUSSION

In this work, we aimed to understand the molecular basis for the growth inhibition of *F. johnsoniae* by *P. koreensis* on a route to elucidating interactions within the rhizosphere microbiome. We have shown that *P. koreensis* inhibits *F. johnsoniae* growth through the production and secretion of novel secondary metabolites, koreenceine A-D (**1–4**), which have structural similarity with the plant metabolite γ-coniceine. Based on the biosynthetic gene cluster identified through our genetic screen, we propose a type II polyketide biosynthetic pathway for these bacterial alkaloids. Traditionally type II polyketide synthases are iterative heterodimer systems. It is currently unclear if the two heterodimer systems present in the biosynthetic cluster act in a modular or iterative manner, but the number of putative β-ketoacyl synthase genes in the pathway is consistent with modular biosynthesis. Plants also use a polyketide pathway mediated by an iterative type III polyketide synthase, providing a new example of convergent evolution between these organisms for the synthesis of related alkaloids. The piperidine core of the koreenceine metabolites is found in well-known plant alkaloids, such as the active cytotoxin γ-coniceine from poison hemlock (*C. maculatum*). Thus, koreenceine alkaloids may play roles in inter-bacterial and inter-Domain communication or inhibition that changes the rhizosphere community structure.

Members of the genus *Pseudomonas* are ubiquitous in nature and thrive in soil, on plants, and on moist surfaces. *P. koreensis* and other members of the *Pseudomonas fluorescens* complex are often studied for their capacity to colonize the rhizosphere and protect plants from pathogens. Previous research demonstrated that *P. fluorescens* suppresses plant disease through production of phenazine-1-carboxylic acid (PCA), which targets the fungal pathogen *Gaeumannomyces graminis* (20). *P. fluorescens* also produces a suite of antimicrobial compounds including 2,4-diacetylphloroglucinol, pyoluteorin, pyrrolnitrin, lipopeptides, and hydrogen cyanide (4), and members of the *P. fluorescens* complex also produce plant hormones, such as indole acetic acid and gibberellic acid, that stimulate plant growth. In this paper, we expand the known repertoire of metabolites from the *P. fluorescens* complex with koreenceine A-D. Unlike most of the *P. fluorescens* metabolites that inhibit fungal pathogens, the koreenceines mediate interactions between *P. koreensis* and diverse members of the Bacteroidetes, including *F. johnsoniae*, in a family-specific manner (6). Competition between members of the *P. fluorescens* complex and *Flavobacterium* spp. in natural settings has been reported; *in vivo* studies showed a selective reduction of *Flavobacterium* spp. in the *Arabidopsis thaliana* rhizosphere when *Pseudomonas* sp. CH267 was added to soil (21). Together, these results highlight the relevance of characterizing bacterial-bacterial interactions in the rhizosphere.

We identified a gene cluster necessary for the production of the koreenceine metabolites. Other koreenceine-like clusters have been predicted to mediate (22) and others are associated with (17, 18) antagonistic activity against diverse microorganisms. Thirty-five percent of *P. koreensis* and *P. mandelii* genomes in NCBI contain this cluster and there are at least 160 *Pseudomonas* genomes harboring a related cluster, indicating the widespread nature of this cluster among *Pseudomonas* spp. Despite its ubiquity, structural characterization of the products of the biosynthetic cluster have been identified slowly.

Bioinformatic analysis of the proposed activities of the genes in the cluster enabled us to propose a biosynthetic pathway for the formation of a C_14_ polyketide with a piperidine-type ring from a non-canonical type II PKS system (Fig. 2). In bacteria, piperidine-type rings could be derived from lysine cyclization (23), as observed in plants, or by a two-step reduction-transamination route of polyketide intermediates (24). We propose that the multidomain protein KecF may direct both steps: reduction of the acyl-intermediate to generate the acyl-aldehyde and transamination. This differs from the established route, in which the two activities are encoded in different genes, and the reduction domain represents the terminal domain of a type I polyketide synthase (Fig. 4) (24). We propose that β-ketoacyl synthase(s) incorporate trans-2-decenoyl-, decanoyl-, or 3-hydroxy-decanoyl units to generate koreenceine A, B or C, respectively. It is unclear if these acyl units are recruited as CoA esters from the β-oxidation pathway or as ACP esters from fatty acid synthesis, although the *R* configuration in koreenceine D suggests substrate sampling from the fatty acid synthesis pool (*i.e*., fatty acyl-ACP).

Koreenceine A-C share structural features with γ-coniceine, the metabolite responsible for the toxicity of the poison hemlock, a plant once used in death sentences, and the means by which Socrates took his own life after receiving such a sentence (399 BC). The structural similarity is the result of convergent evolution. Their functions may also be related—γ-coniceine and its derivative make hemlock *(Conium maculatum*) toxic to animals (9) and *P. koreensis* might protect plant roots with koreenceines. γ-coniceine is also considered a plant hormone (9, 25). Thus, future work will focus on characterizing the effect of koreenceine A-C on plant development and protection.

## ACKNOWLEGMENTS

We gratefully acknowledge Dr. Sailendharan Sudakaran for discussing phylogenetic analysis. We thank Dr. Hans Wildschutte for *Pseudomonas* sp SWI36 and its mutant. This work was supported by the Office of the Provost at Yale University, funding from the Wisconsin Alumni Research Foundation through the University of Wisconsin–Madison Office of the Vice Chancellor for Research and Graduate Education, and NSF grant MCB-1243671.

## MATERIALS AND METHODS

### Bacterial strains and culture conditions

*F. johnsoniae CI04, P. koreensis CI12, B. cereus* UW85, *Pseudomonas* sp. SWI36, *Flavobacterium johnsoniae* CI64, *Chryseobacterium* sp. CI02, *Chryseobacterium* sp. CI26, *Sphingobacterium* sp. CI01, and *Sphingobacterium* sp. CI48 were propagated on 1/10th-strength tryptic soy agar and grown in liquid culture in ½-strength tryptic soy broth (TSB) at 28°C with vigorous shaking.

### Production of root exudates and defined media

Soybean seeds were surface sterilized with 6% sodium hypochlorite for 10 min, washed with sterile deionized water, transferred to water agar plates, and allowed to germinate for three days in the dark at 25°C. Seedlings were grown in a hydroponic system using modified Hoagland’s plant growth solution (26), which was collected after 10 days of plant growth in an environmental chamber (12-h photoperiod, 25°C), filter sterilized and stored at −20°C until used as root exudate. A defined medium was based on basal salt medium (1.77 g mL^-1^ Na_2_HPO_4_; 1.70 g mL^-1^ KH_2_PO_4_; 1.00 g mL^-1^ (NH_4_)_2_SO_4_; 0.16 g mL^-1^ MgCl_2_*6H_2_O; 5.00 g mL^-1^ FeSO_4_*7H_2_O). A carbon source (pyruvate, mannitol, or glucose) was added to a final concentration of 4 mM. An amino acid mix of equal parts alanine, aspartate, leucine, serine, threonine, and valine was added to the root exudate or the defined minimal media at a final concentration of 6 mM. Individual amino acids and *N*-methyl-DL-aspartate were also added to a final concentration of 6 mM.

### *P. koreensis* mutant library generation by transposon mutagenesis

*P. koreensis* CI12 and *E. coli* S17–1*λ*pir with pSAM_BT20 (27) with ampicillin (100 μg mL^-1^) were first grown individually for 16 h in LB at 28°C and 37°C, respectively, with agitation. Cells were washed and resuspended in LB to an OD_600_ = 2.0. One volume of *E. coli* S17–1*λ*pir with pSAM_BT20 was mixed with two volumes of *P. koreensis* CI12 were mixed. Cells were harvested (6000 × g, 6 min), resuspended in 100 μL of LB, and spotted on LBA. Plates were incubated at 28°C for 16 h. Each conjugation mixture was scraped off the plate, resuspended in 2.5 mL of LB, and 350-μL aliquots were plated on LB containing gentamicin (50 μg mL^-1^) and chloramphenicol (10 μg mL^-1^) to select for *P. koreensis* CI12 transconjugants. Plates were incubated for two days at 28°C.

### Genetic screen of *P. koreensis* mutants defective in inhibitory activity

*P. koreensis* CI12 mutants were grown for 16 h in 96-deepwell plates filled with ½-strength TSB, covered with sterile breathable sealing films, and incubated at 28°C with agitation. For each plate, the first well was inoculated with wild type *P. koreensis* CI12, and the last well was left without *P. koreensis* CI12. *F. johnsoniae* CI04 was grown and washed as described above. Root exudate was inoculated with ~10^7^ *F. johnsoniae* CI04 cells per mL, and 200 μL aliquots were added to each well of 96-well microplates. Next, two μL from each mutant *P. koreensis* CI12 culture was transferred to the corresponding wells on the microplates, which were then covered by sealing films and incubated at 28°C with slight agitation for two days. Five μL from each well were then spotted on Casitone-yeast extract agar (CYE) (10 g L^-1^ casitone; 5 g L^-1^ yeast extract; 8 mM MgSO4; 10 mM Tris buffer; 15 g L^-1^ agar) containing kanamycin (10 μg mL^-1^) to select for *F. johnsoniae*, and plates were incubated at 28°C for two days. Mutants that did not inhibit *F. johnsoniae* were streaked on a second plate for further analysis. The loss of inhibitory activity of candidate *P. koreensis* mutants was verified in a second co-culture, and mutant growth was then compared to wild-type growth to rule out candidates that failed to inhibit *F. johnsoniae* due to their own growth deficiency.

### Location transposons in *P. koreensis* mutants defective in *F. johnsoniae* inhibition

For each mutant, one mL of liquid culture grown for 16 h was harvested (6000 × g, 6 min), and cells were resuspended in 400 mL of TE (10 μM TrisHCl pH 7.4; 1 μM EDTA pH 8.0). Samples were boiled for 6 min, centrifuged (6000 × *g*, 6 min), and two μL of supernatant was used as a template for DNA amplification. Transposon locations were determined by arbitrarily primed PCR which consisted of a nested PCR using first-round primer GenPATseq1 and either AR1A or AR1B and second-round primer GenPATseq2 and AR2 (Table 2). PCR products from the second round were purified by gel extraction (QIAquick Gel Extraction Kit; QIAGEN) and then sequenced using primer GenPATseq2.

**Table 2.**
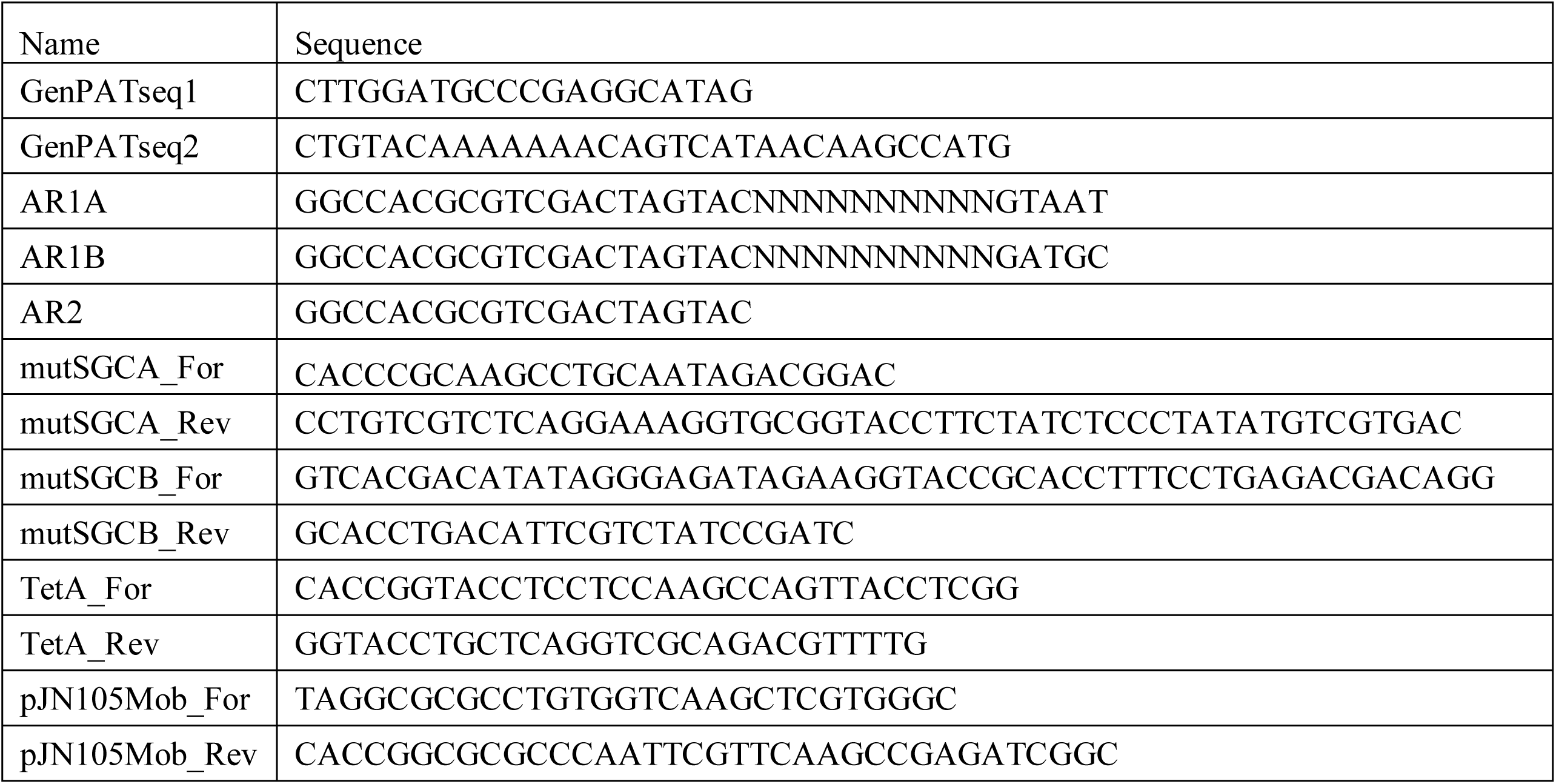
Primers used in this study.

### Chromosomal deletion of the koreenceine gene cluster in *P. koreensis*

The koreenceine cluster was deleted by allelic exchange and replaced with a tetracycline resistant cassette. The *kecA-K* deletion cassette was constructed by a modified version of overlap extension (OE) PCR strategy. Fragments one kb upstream and one kb downstream of the *kecA-K* genes were amplified using primers mutSGCA_For/mutSGCA_Rev and mutSGCB_For/mutSGCB_Rev respectively (Table 2). The PCR products were cloned in pENTR/D-TOPO, generating pkecA-K_ENTR. Primers mutSGCA_Rev and mutSGCB_For were designed to include a KpnI site in their overlapping region to allow introduction of a resistance gene. A tetracycline resistance cassette was amplified from pACYC184 using primers TetA_For/TetA_Rev, which contain KpnI sites in the 5’ region, and cloned into pENTR/D-TOPO to generate pTetA_ENTR. A *mob* element was amplified from pJN105 using primers pJN105Mob_For/pJN105Mob_Rev (Table 2), in which an AscI site in the 5’ region was added, and cloned in pENTR/D-TOPO, generating pmob_ENTR. The tetracycline cassette was recovered from pTetA_ENTR using KpnI, and cloned between the region upstream and downstream of the pkecA-K_ENTR, and the *mob* element was recovered from pmob_ENTR using AscI, and cloned into an AscI site in the pENTR backbone, generating pkecA-K_TetA_mob_ENTR. Conjugation mixtures of *P. koreensis* CI12 and *E. coli* S17–1λpir carrying the pkecA-K_TetA_mob_ENTR vector were prepared following the procedure for the transposon mutant generation. Double recombinant *P. koreensis* CI12 transconjugants were selected by their ability to grow on tetracycline (10 μg mL^-1^) and inability to grow on kanamycin (50 μg mL^-1^). The *kekA-K* deletion mutant was confirmed by PCR using primers mutSGCA_For and mutSGCB_Rev. The *kekA-K* deletion mutant was further confirmed by evaluating growth of *F. johnsoniae*, and other members of the Bacteroidetes, in its presence.

### General information for the analysis and identification of metabolites

^1^H and 2D-(gCOSY, gHSQC, and gHMBC) NMR spectra were obtained on an Agilent (USA) 600 MHz NMR spectrometer with a cold probe, and the chemical shifts were recorded as *δ* values (ppm) with methanol-*d*_4_ as the standard NMR solvent. Materials were routinely analyzed on an Agilent 6120 single quadrupole liquid chromatography-mass spectrometry (LC/MS) system (Column: Phenomenex kinetex C_18_ column, 250 × 4.6 mm, 5 μm; Flow rate: 0.7 ml min^-1^; Mobile phase composition: H_2_O and acetonitrile (ACN) containing 0.1% trifluoroacetic acid (TFA); Method: 0–30 min, 10–100% ACN; hold for 5 min, 100% ACN; 1 min, 100–10% ACN). High-resolution electrospray ionization mass spectrometry (HR-ESIMS) data were obtained using an Agilent iFunnel 6550 Q-TOF (quadrupole-time-of-flight) mass spectrometer fitted with an electrospray ionization (ESI) source coupled to an Agilent (USA) 1290 Infinity high performance liquid chromatography (HPLC) system. Open column chromatography was carried out on a Waters Sep-Pak^®^ Vac 35cc (10g) C_18_ column. Metabolite isolations were performed using an Agilent (USA) Prepstar HPLC system with an Agilent (USA) Polaris C_18_-A 5 μm (21.2 × 250 mm) column, a Phenomenex (USA) Luna C_18_(2) (100Å) 10 μm (10.0 × 250 mm) column, a Phenomenex (USA) Luna C_8_(2) (100Å) 10 μm (10.0 × 250 mm) column, and an Agilent Polaris 5 Amide-C_18_ (250 × 10.0 mm) column.

### Isolation of metabolites

*P. koreensis CI12* was grown in defined medium with pyruvate as carbon source and supplemented with the amino acid mix or 3mM of glutamate for 3 days. Crude extract was generated by liquid-liquid extraction using one volume of 2-butanol per one volume of filter supernatant, and dried by rotary evaporation. The crude extracts (400 mg) from the 12-L culture supernatant were resuspended in water and methanol (1:1 ratio), adsorbed onto Celite^®^110, and dried by rotary evaporation. The resulting powdery materials were loaded on the Waters Sep-Pak^®^ Vac 35cc (10g) C_18_ cartridge, and the metabolites were separated by solvent fractionation, eluting with a step gradient from 20–100% aqueous methanol to yield five sub-fractions (20%, 40%, 60%, 80%, and 100% methanol containing 0.1% TFA). Reversed-phase LC-MS analysis (10–100% aqueous acetonitrile in 0.1% trifluoroacetic acid, 30-min gradient) revealed that the 60% fraction included both molecules **1** and **2**, and the fraction was dried under reduced pressure. This fraction (60 mg) was then separated by reversed-phase HPLC equipped with an Agilent Polaris C_18_-A 5 μm (21.2 × 250 mm) column with an isocratic solvent system (50% acetonitrile in water, 0.1% TFA, over 20 min, 8 mL min^-1^ 1-min fraction collection interval). Compound **1** from the pooled fraction (11+12) (*t*_R_ = 25.3 min, 0.2 mg) was partially purified over the Phenomenex Luna C_8_ (2) 10 μm (10.0 × 250mm) column with a linear gradient elution (20–80% acetonitrile in water, 0.1% TFA, over 30 min). The combined HPLC fraction (11+12) was subsequently purified by reversed-phase HPLC (Phenomenex Luna C_18_ (2) 10 μm (10.0 × 250mm) column) with a linear gradient elution (20–80% acetonitrile in water, 0.1% TFA, over 30 min) to yield pure compound **2** (*t*_R_ = 25.8 min, 1.2 mg). Compound **4** was detected in the 80% aqueous methanol Sep-Pak fraction and was separated over an Agilent Polaris C_18_-A 5 μm (21.2 × 250 mm) column (Flow rate: 8.0 ml/min; Gradient elution: 10–100% aqueous acetonitrile in 0.1% TFA for 30 min, 1-min fraction collection). HPLC fraction 24 was then separated over the Phenomenex Luna C_18_ (2) 10 μm (10.0 × 250mm) column with 50–100% acetonitrile in water containing 0.1% TFA over 30 min at a flow rate of 4 ml min^-1^ followed by the subjection to Agilent Polaris 5 Amide-C_18_ (250 × 10.0 mm) with the same elution system (Flow rate: 4 ml min^-1^: Purification method: 50–100% acetonitrile in water containing 0.1% TFA over 30 min) to yield pure compound **4** (*t*_R_ = 9.43 min, 0.7 mg).

(*E*)-6-(non-1-en-1-yl)-2,3,4,5-tetrahydropyridine (**1**): colorless solid; ^1^H NMR (CD_3_OD, 600 MHz) *δ* 7.21–7.12 (1H, m, H-8), 6.35 (1H, d, *J* = 16.0 Hz, H-7), 3.59 (2H, m, H-2), 2.91 (2H, m, H-5), 2.30 (2H, m, H-9), 1.77 (2H, m, H-3), 1.71 (2H, m, H-4), 1.42 (2H, m, H-10), 1.27–1.20 (8H, m, H-11, H-12, H-13, H-14), 0.84 (3H, t, *J* = 7.0 Hz, H-15); HR-ESI-QTOF-MS [M+H]^+^ *m/z* 208.2067 (calcd for C_14_H_26_N, 208.2065).

6-nonyl-2,3,4,5-tetrahydropyridine (**2**): colorless solid; ^1^H NMR (CD_3_OD, 600 MHz) *δ* 3.53 (2H, t, *J* = 5.5 Hz, H-2), 2.72 (2H, t, *J* = 6.1 Hz, H-5), 2.53 (2H, m, H-7), 1.73 (2H, m, H-3), 1.68 (2H, m, H-4), 1.54 (2H, dt, *J* = 14.6, 6.8 Hz, H-8), 1.28–1.15 (12H, m, H-9, H-10, H-11, H-12, H-13 H-14), 0.83 (3H, t, *J* = 7.0 Hz, H-15), ^13^C NMR (CD_3_OD, 125 MHz) *δ* 192.1 (C-6),44.4 (C-2), 37.7 (C-7), 31.6 (C-13), 29.4 (C-5), 28.0–29.0 (C-9, C-10, C-11, C-12), 25.5 (C-8), 22.5 (C-14), 19.2 (C-3), 16.8 (C-4), 14.4 (C-15); HR-ESI-QTOF-MS [M+H]^+^ *m/z* 210.2224 (calcd for C_14_H_28_N, 210.2222).

(*R*)-*N*-(4-chlorobutyl)-3-hydroxydecanamide (**4**): colorless solid; ^1^H NMR (CD_3_OD, 600 MHz) *δ* 3.94 (1H, m, H-3), 3.59 (2H, t, *J* = 6.5 Hz, H-4′), 3.20 (2H, m, H-1′), 2.27 (2H, m, H-2), 1.78 (2H, m, H-3′), 1.63 (2H, dt, *J* = 14.3, 7.0 Hz, H-2′), 1.43 (2H, m, H-4), 1.36–1.25 (10H, m, H-5, H-6, H-7, H-8, H-9), 0.90 (3H, t, *J* = 6.9 Hz, H-10), ^13^C NMR (CD_3_OD, 125 MHz) *δ* 172.6 (C-1), 68.2 (C-3), 44.1 (C-4′), 43.8 (C-2), 37.9 (C-1′), 36.8 (C-4), 31.8 (C-8), 29.5 (C-3′), 29.4 (C-5 or C-6, C-7), 26.3 (C-2′), 22.6 (C-5 or C-6, C-9), 13.4 (C-10); HR-ESI-QTOF-MS [M+H]^+^ *m/z* 278.1885 (calcd for C_14_H_29_ClNO_2_, 278.1887).

### Determination of absolute configuration of metabolite 4

The absolute configuration of **4** was determined using the modified Mosher’s method with R- and S-a-methoxy-(trifluoromethyl)phenylacetyl chloride (MTPA-Cl) (28). Compound **4** (0.5 mg) was prepared in two vials (0.25 mg), and each sample was dissolved in 250 *μ*L of dried pyridine-*d*_5_ in vials purged with N_2_ gas. Dimethylaminopyridine (DMAP) (0.5 mg) was added to both vials followed by the addition of 5 *μ*L of *S*- and *R*-MTPA-Cl solution (2% v/v) at room temperature. After 18 h, the reaction mixtures were dried under reduced pressure. ^1^H NMR spectra of the Mosher esters (*S*-MTPA ester and *R*-MTPA ester) were collected in methanol-*d*_4_, and the chemical shift differences of the Mosher esters of **4** were calculated in Δ*δ*_S-R_.

### Characterization of *Pseudomonas* sp. SWI36

*Pseudomonas* sp. SWI36 and *P. koreensis* inhibitory interactions against *B. cereus* and *F. johnsoniae* were evaluated with a modified spread-patch method. Strains were grown separately for 20 h. One-mL aliquots of cultures of each strain were centrifuged (6000 × *g*, 6 min), resuspended in one ml of the same medium (undiluted cultures), and a 1:100 dilution of *B. cereus* and *F. johnsoniae* was prepared in the same medium (diluted culture). Nutrient agar plates were spread with 100 of either *B. cereus* or *F. johnsoniae* diluted cultures and spotted with 10 μL of the undiluted cultures of *Pseudomonas* sp. SWI36 and *P. koreensis*. Plates were then incubated at 28°C and inspected for zones of inhibition after two days. Crude extract of *Pseudomonas* sp. SWI36 and *Pseudomonas* sp. SWI36 kecF::Tn culture in nutrient broth were prepared as above. Extracted materials were analyzed on a LC/MS system consisting of a Thermo Fisher Scientific (Waltham, MA) Q Exactive orbitrap mass spectrometer with an electrospray ionization (ESI) source coupled to a Vanquish UHPLC (Column: Thermo Accucore Vanquish C_18_ column, 100 × 2.1 mm, 1.5 μm; Flow rate: 0.2 ml min^-1^; mobile phase composition: H_2_O and acetonitrile (ACN) containing 0.1% trifluoroacetic acid (TFA); method: 0–1 min, 10% ACN; **1–4** min, 10–35% ACN; 4–12 min, 35–70% ACN; 12–16 min, 70–98% ACN; 16–20 min hold with 98% ACN; 20–21 min, 98–10% ACN; 21–23 min, 10% ACN). MS1 scans were acquired with positive ionization over a *m/z* range of 188–1275 with settings of 1e6 AGC, 100 ms maximum integration time, and 70k resolution.

### Phylogenetic analysis

Genetic regions with homology to the koreenceine biosynthetic cluster were identified by BLAST alignment tools (29) using *P. koreensis* CI12 KecF protein sequence in the NCBI database. All the KecF homologues identified were part of koreenceine-like cluster. Genomes harboring koreenceine-like clusters are listed in Table S1. Protein and nucleotide sequence alignments of *kecF* were performed with MAFFT version 7 (30) and were manually adjusted using as a guide the residues-wise confidence scores generated by GUIDANCE2 (31). Best-fit models of amino acid or nucleotide replacement were selected. Evolutionary analyses were inferred by Maximum Likelihood (ML) methods conducted in MEGA X (32). The *P. koreensis* and *P. mandelii* phylogenomic reconstruction was done by the phylogenetic and molecular evolutionary (PhaME) analysis software (33). PhaME identified SNPs from the core genome alignments, and the phylogenetic relationships were inferred by ML using FastTree. Phylogenetic trees were visualized using interactive tree of life (iTOL) (34).

---

**FIG S1.**
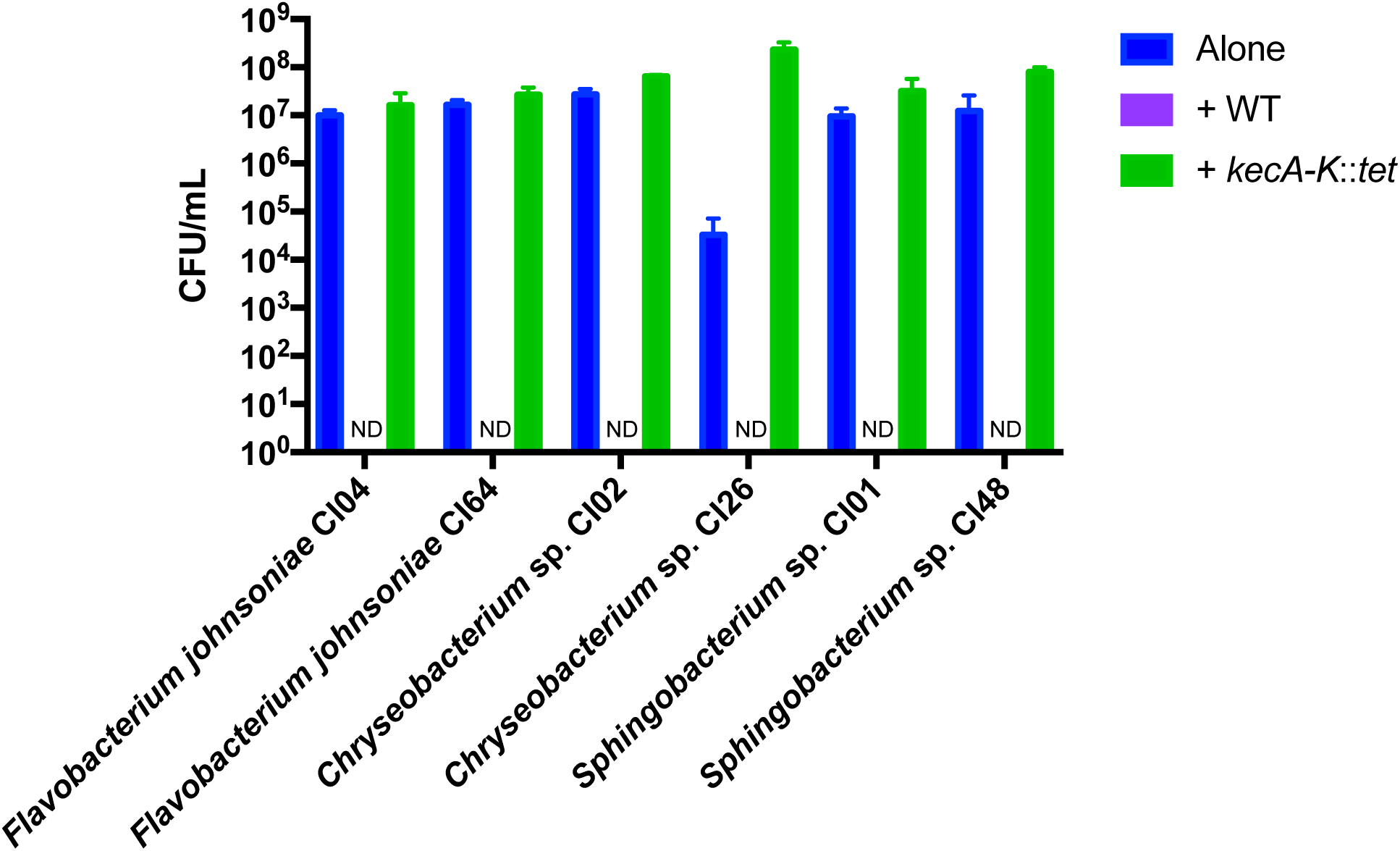
Genetic validation of the role of koreenceine gene cluster in inhibition of *F. johnsoniae* and other Bacteroidetes. Several Bacteroidetes members of the soybean rhizosphere population when grown alone, with *P. koreensis* wild type, or *P. koreensis* CI12 *kecA-K*::*tet*. ND, not detected.

**FIG S2.**
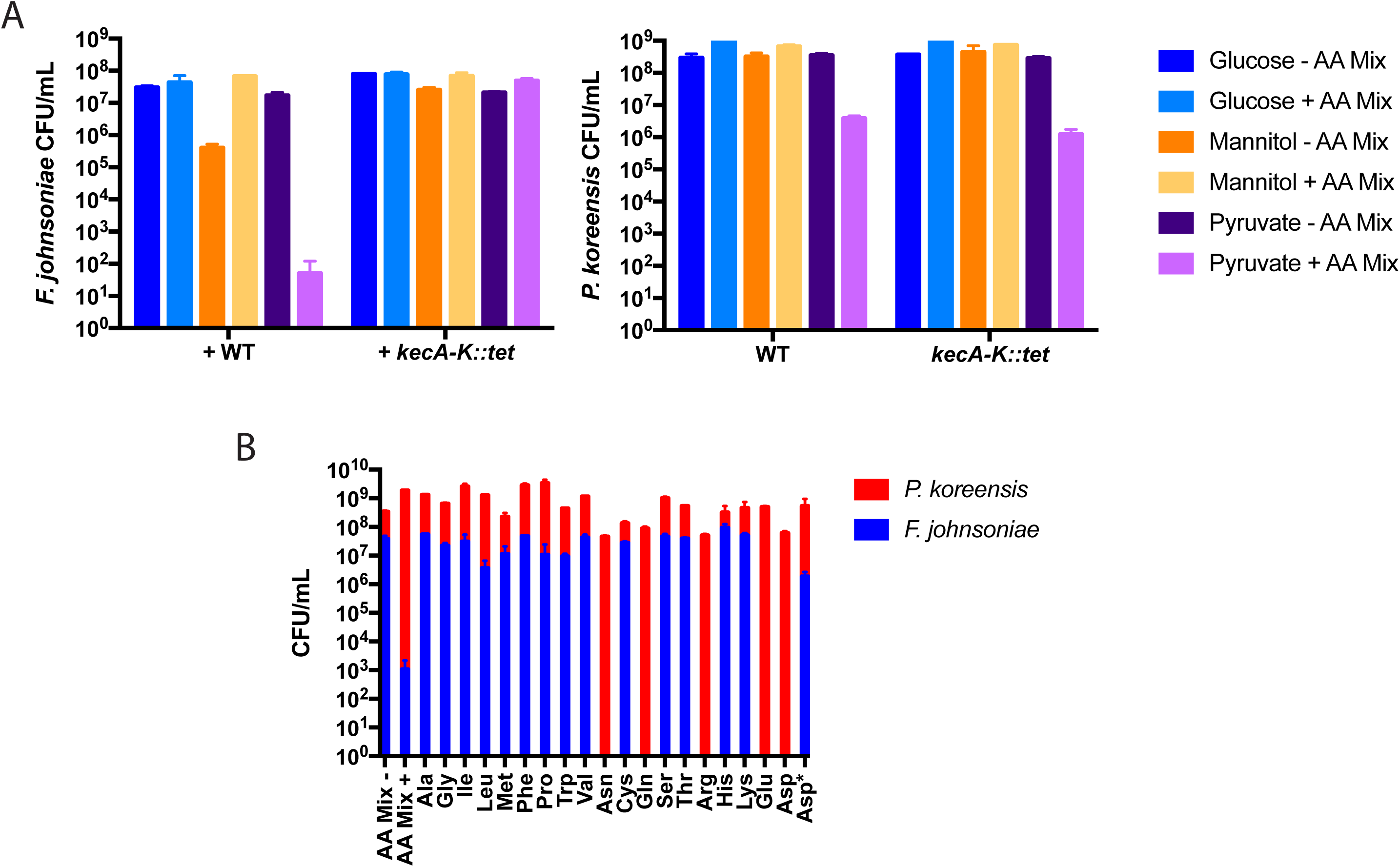
Development of a minimal medium to mimic *P. koreensis* inhibition against *F. johnsoniae*. A) *P. koreensis* and *F. johnsoniae* in co-culture in a define medium with glucose, mannitol or pyruvate as a carbon sources with or without amino acid mixture. B) *P. koreensis* and *F. johnsoniae* in co-culture in a defined minimal media with or without amino acid mixture, or a single amino acid, or N-methyl-DL-aspartate (Asp*) a non-hydrolysable analog of aspartate.

**FIG S3.**
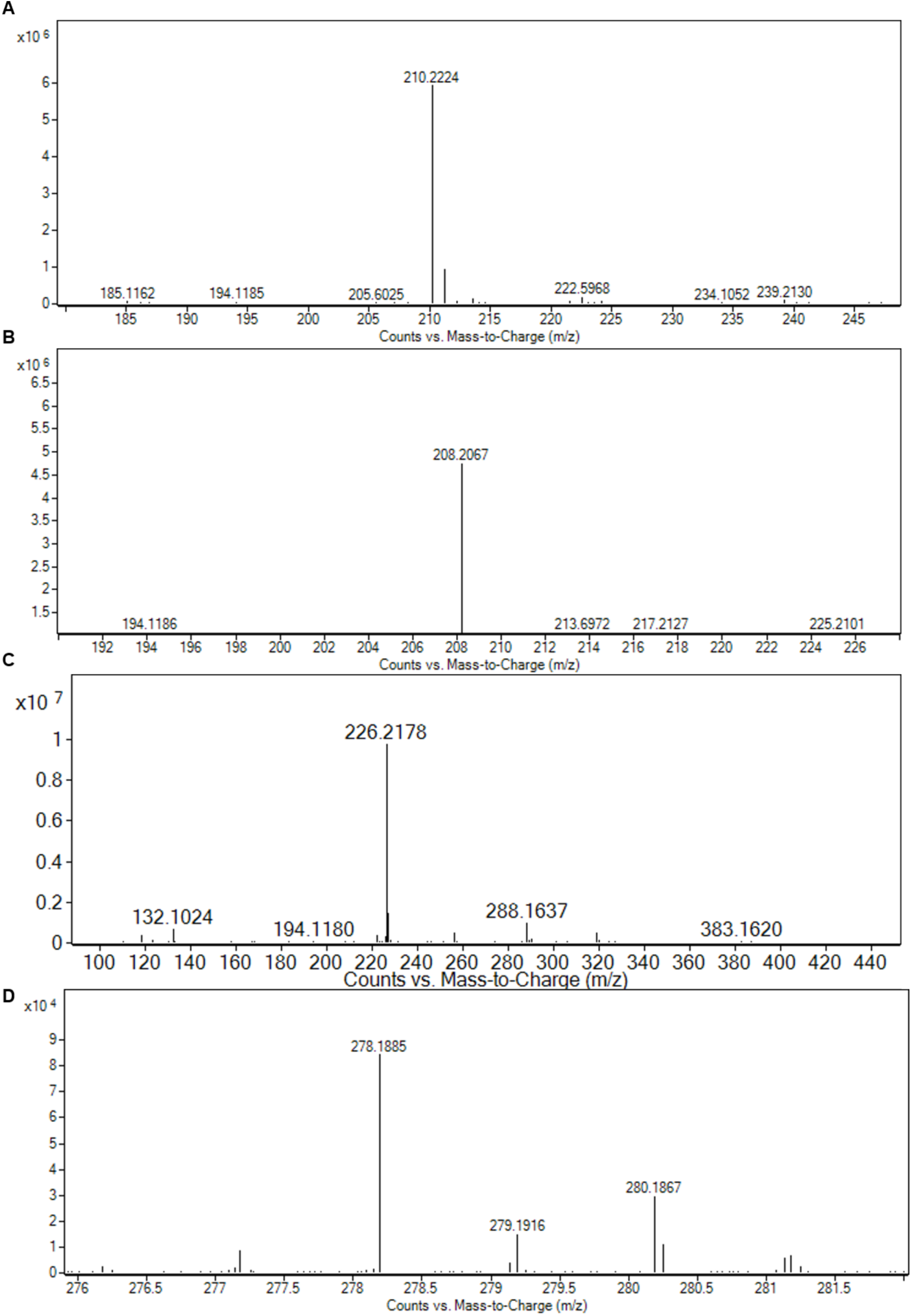
HRESIQTOFMS spectral data of compounds **1–4**

**FIG S4.**
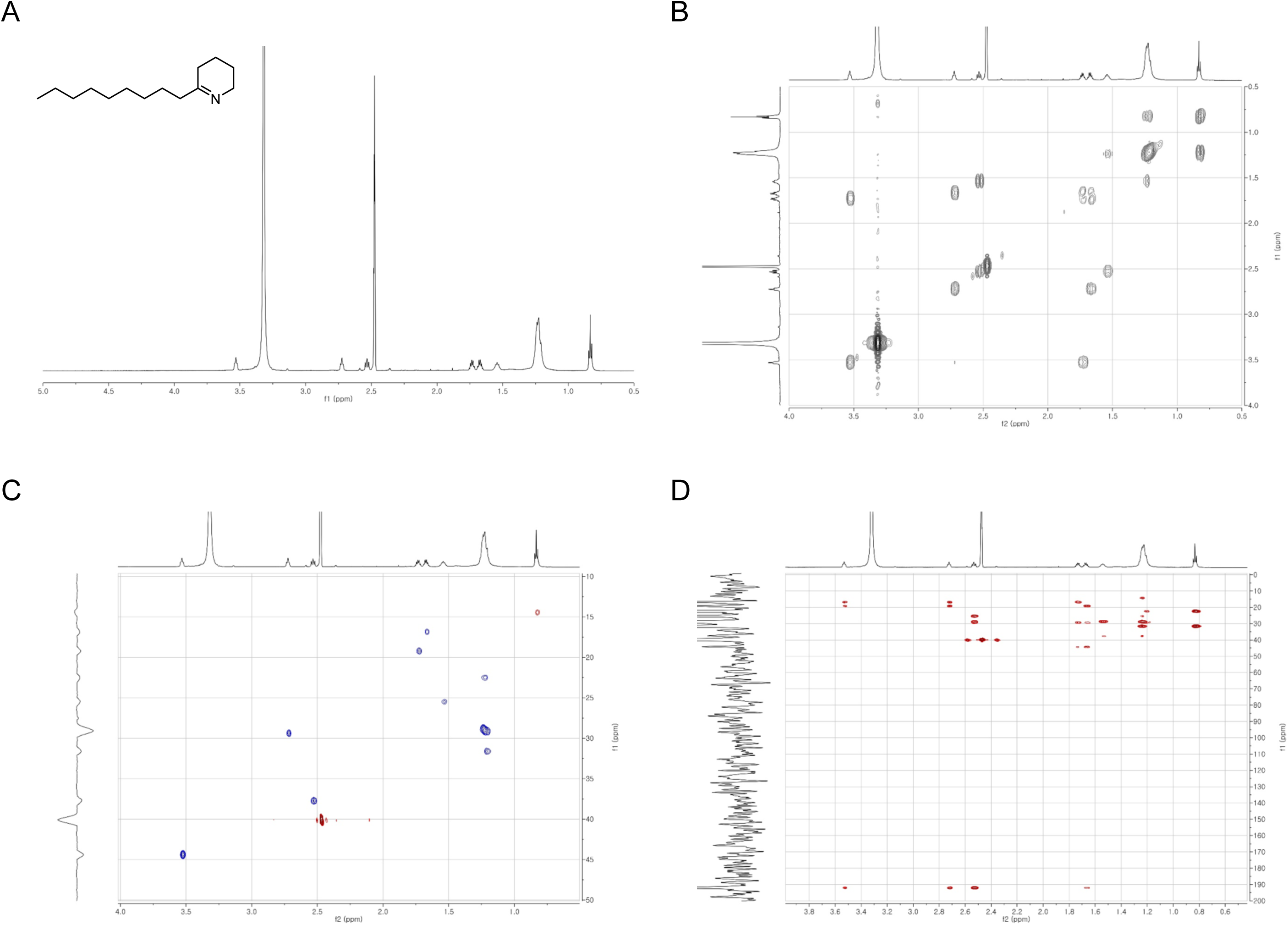
NMR spectra of compound **1**

**FIG S5.**
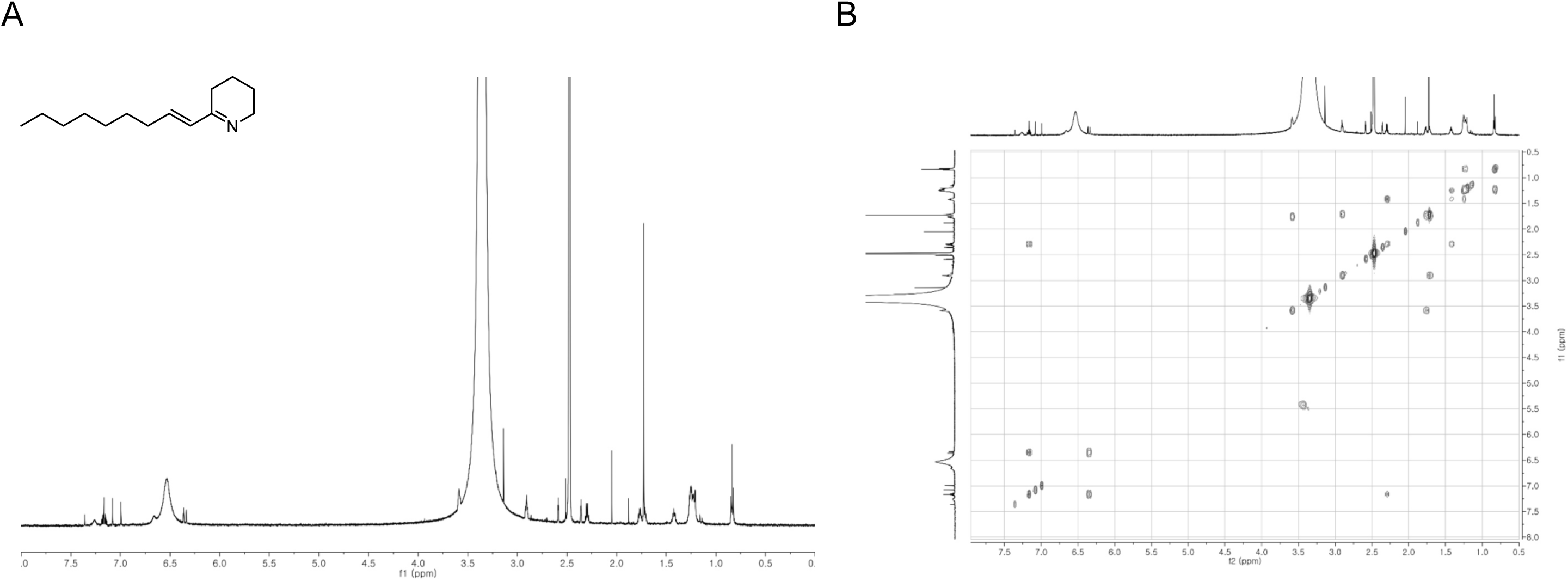
NMR spectra of compound **2**

**FIG S6.**
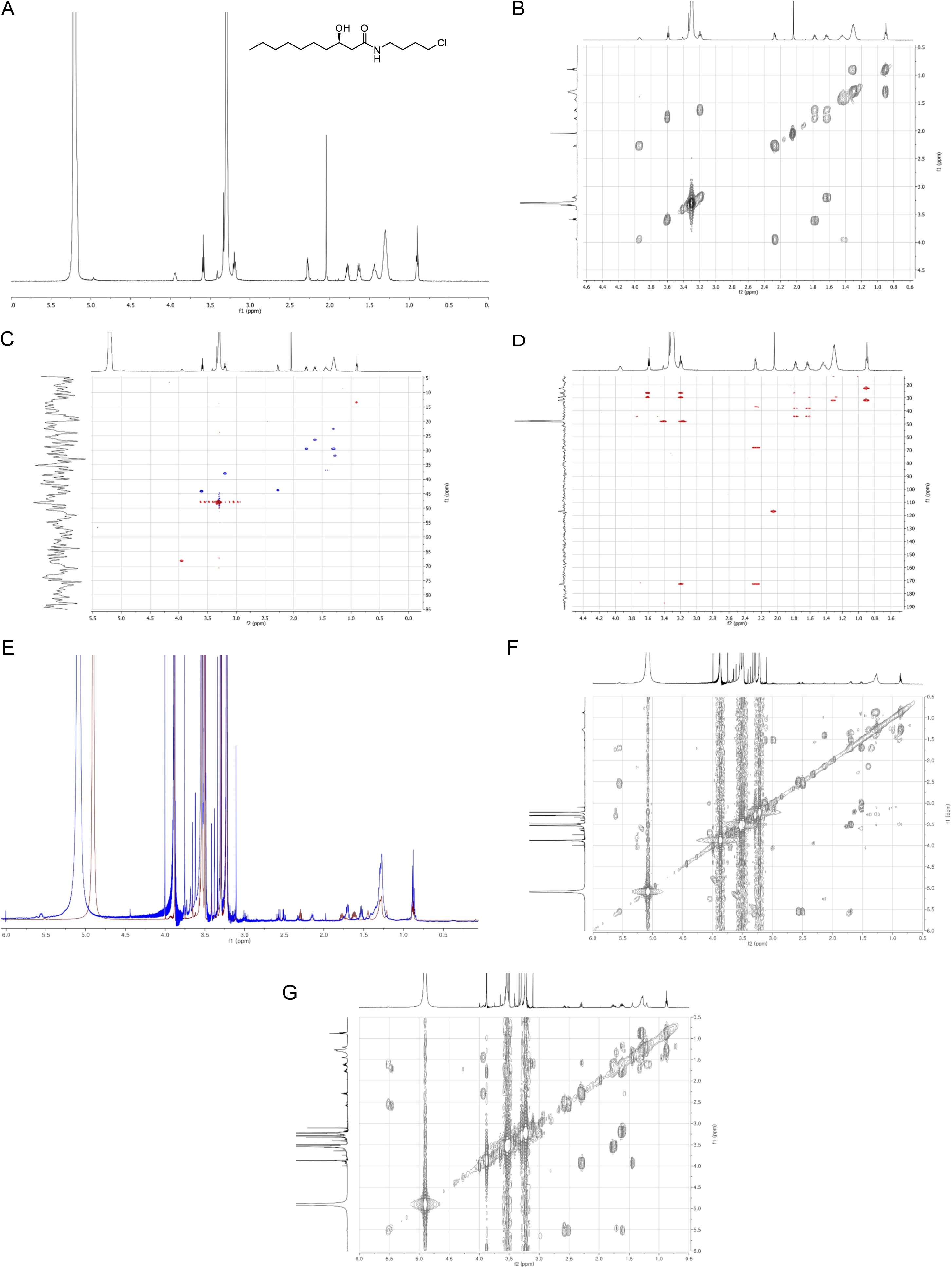
(A-D) NMR spectra of compound **4** (E) ^1^H NMR spectra comparison of *S*-MTPA (blue) and *R*-MTPA (maroon) of compound **4** (F) gCOSY NMR spectrum of *S*-MTPA of compound **4** (G) gCOSY NMR spectrum of *R*-MTPA of compound **4**

**FIG S7.**
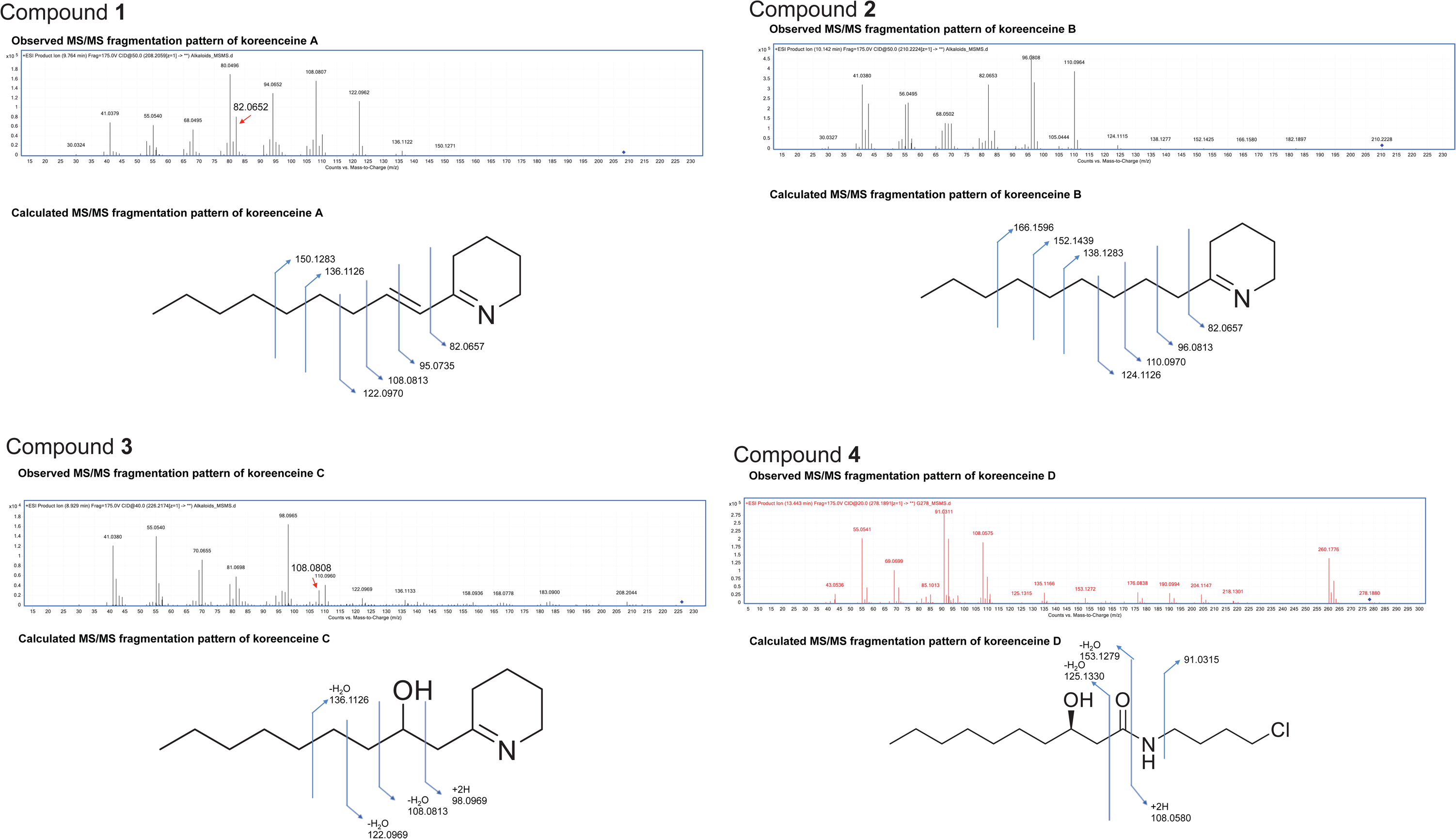
Tandem MS spectra of compounds **1–4**

**FIG S8.**
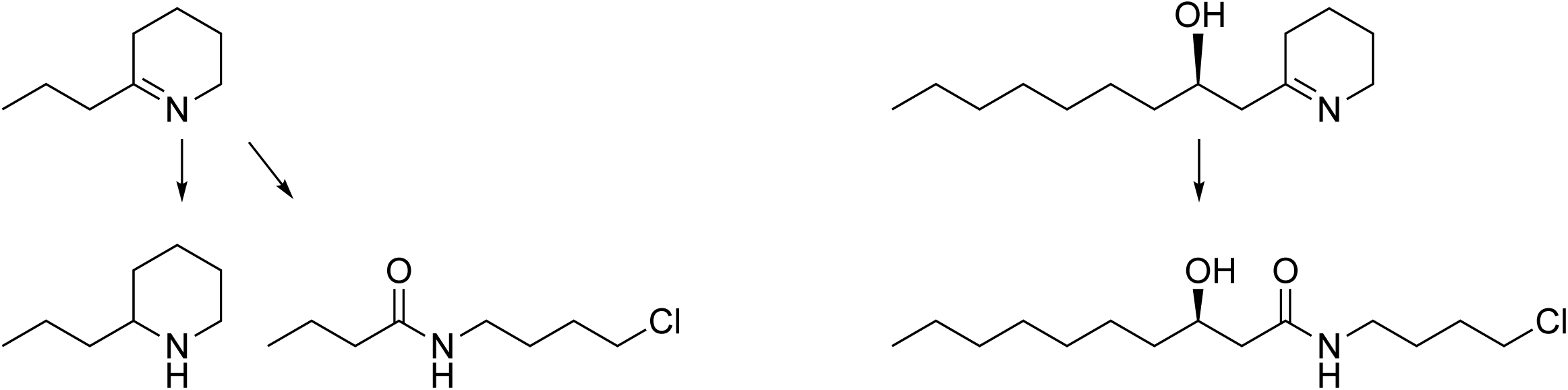
Predicted uncharacterized halogenation reaction and oxidation/reduction of γ-coniceine and koreenceine C.

**FIG S9.**
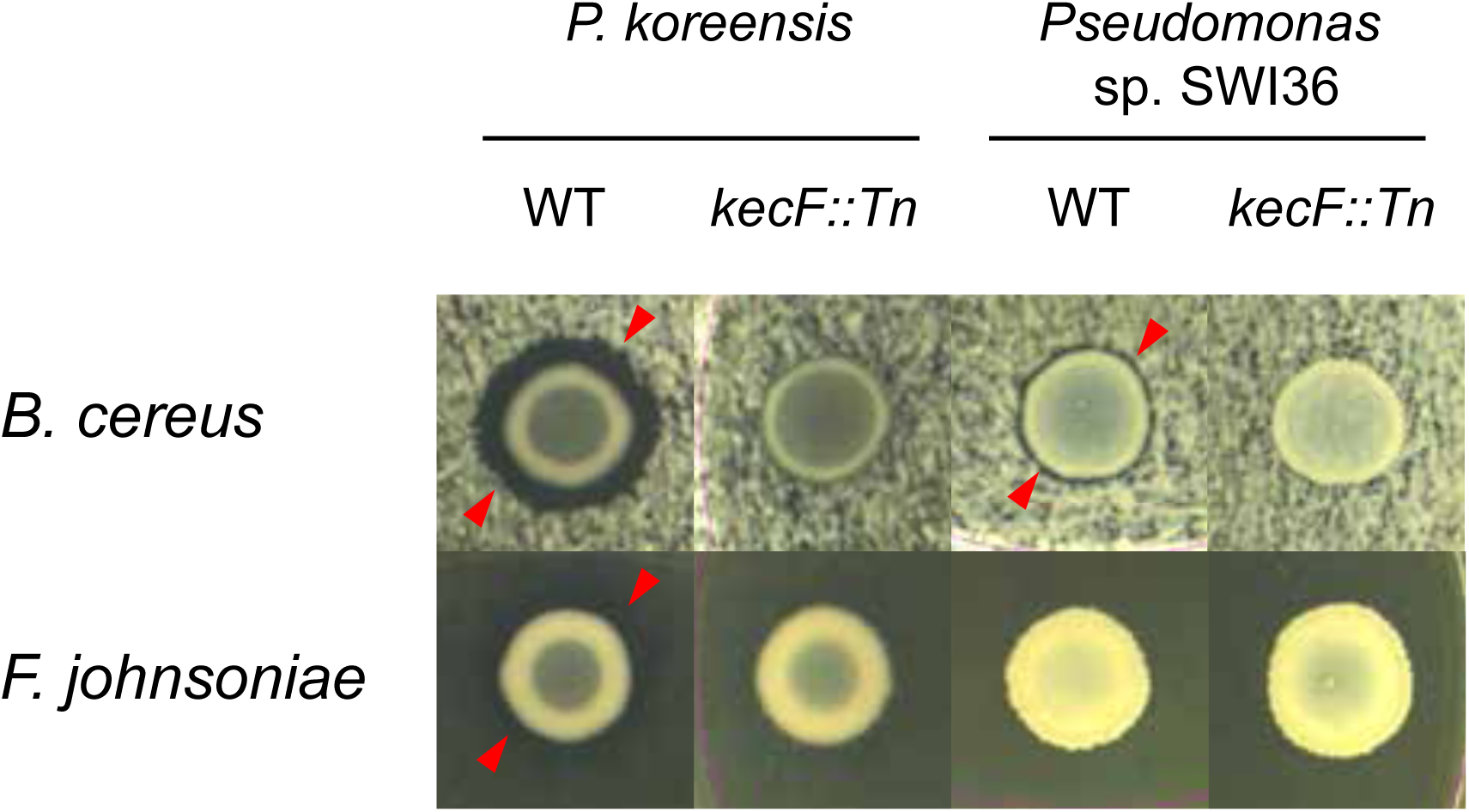
Activity profile of *P. koreensis* and *Pseudomonas* sp. SWI36 against *B. cereus* and *F. johnsoniae*.

**FIG S10.**
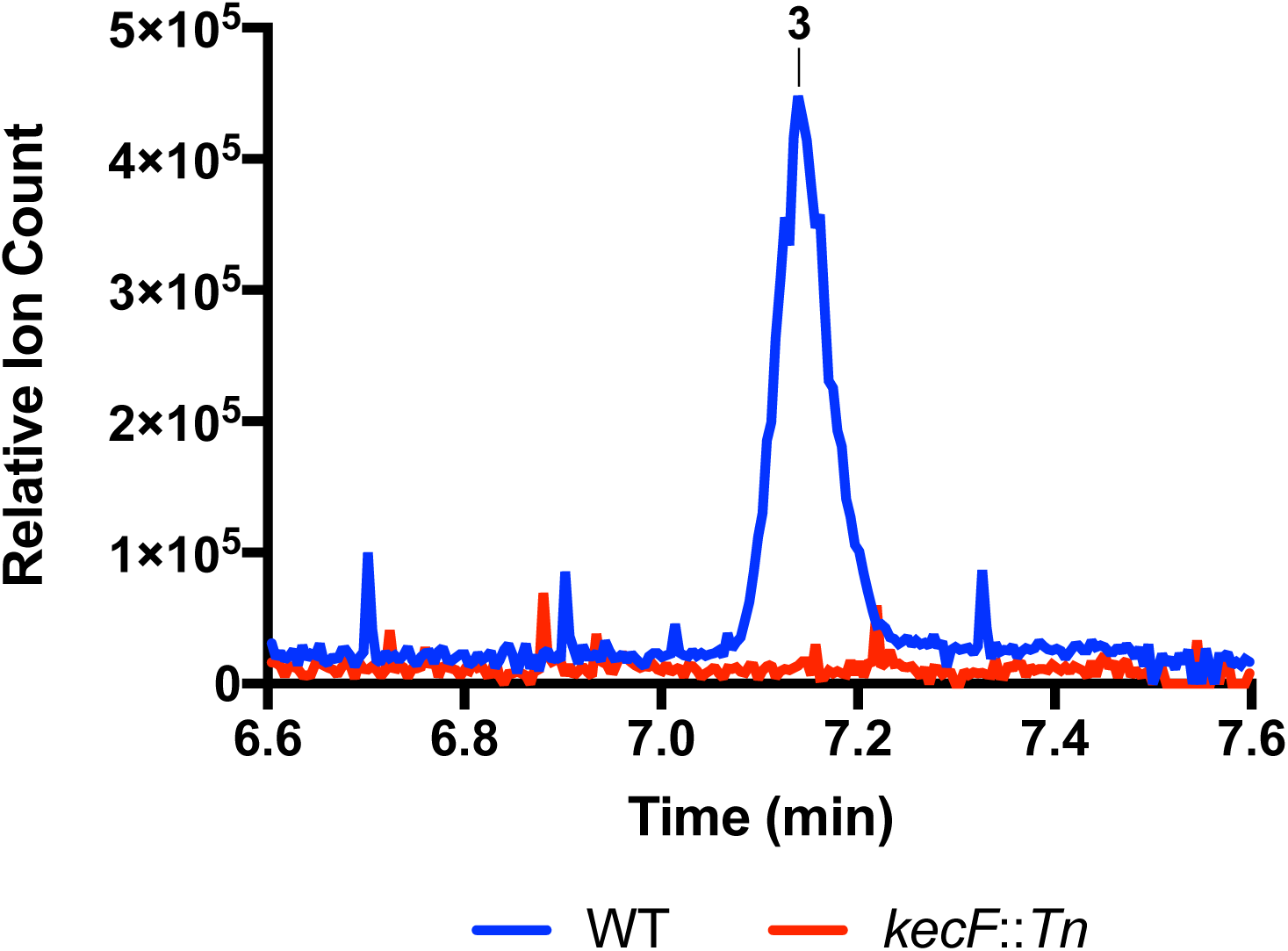
Extracted ion chromatograms of koreenceine C for *Pseudomonas* sp SWI36 wild type and *kecF*::*Tn* mutant.

**Table S1.** Bacterial genomes containing gene clusters with high similarity to the koreenceine cluster.

| Name | Abbreviation | GenBank assembly accession | Koreenceine-like cluster clade |
| --- | --- | --- | --- |
| [*Flavobacterium*] sp. 29 | 29 | GCA_002754355 | A |
| *Pseudomonas asplenii* 4A7 | 4A7 | GCA_002891515 | A |
| *Pseudomonas baetica* a390 | a390 | GCA_003031005 | A |
| *Pseudomonas baetica* LMG 25716 | LMG25716 | GCA_002813455 | A |
| *Pseudomonas fluorescens* NZ011 | NZ011 | GCA_000276585 | A |
| *Pseudomonas fluorescens* BW11P2 | BW11P2 | GCA_001679645 | A |
| *Pseudomonas fluorescens* C3 | C3 | GCA_000967955 | A |
| *Pseudomonas fluorescens* H16 | H16 | GCA_000802985 | A |
| *Pseudomonas fluorescens* MS82 | MS82 | GCA_003055645 | A |
| *Pseudomonas jessenii* LBp-160603 | LBp160603 | GCA_003205375 | A |
| *Pseudomonas koreensis* CI12 | CI12 | GCA_002003425 | A |
| *Pseudomonas mandelii* JR-1 | JR1 | GCA_000257545 | A |
| *Pseudomonas mandelii* PD30 | PD30 | GCA_000690555 | A |
| *Pseudomonas mandelii* LMG 21607 | LMG21607 | GCA_900106065 | A |
| *Pseudomonas prosekii* LMG 26867 | LMG26867 | GCA_900105155 | A |
| *Pseudomonas prosekii* P2406 | P2406 | GCA_003122265 | A |
| *Pseudomonas prosekii* P2673 | P2673 | GCA_003122305 | A |
| *Pseudomonas* sp. 31-12 | 3112 | GCA_003151075 | A |
| *Pseudomonas* sp. 43NM1 | 43NM1 | GCA_002836905 | A |
| *Pseudomonas* sp. AD21 | AD21 | GCA_002878485 | A |
| *Pseudomonas* sp. B15(2017) | B152017 | GCA_002113215 | A |
| *Pseudomonas* sp. B17(2017) | B172017 | GCA_002113765 | A |
| *Pseudomonas* sp. B2(2017) | B22017 | GCA_002113685 | A |
| *Pseudomonas* sp. B20(2017) | B202017 | GCA_002113125 | A |
| *Pseudomonas* sp. B26(2017) | B262017 | GCA_002113045 | A |
| *Pseudomonas* sp. B35(2017) | B352017 | GCA_002113645 | A |
| *Pseudomonas* sp. B7(2017) | B72017 | GCA_002112745 | A |
| *Pseudomonas* sp. FSL W5-0299 | FSLW50299 | GCA_002005125 | A |
| *Pseudomonas* sp. FW305-BF8 | FW305BF8 | GCA_002883215 | A |
| *Pseudomonas* sp. GM16 | GM16 | GCA_000282155 | A |
| *Pseudomonas* sp. GM24 | GM24 | GCA_000282235 | A |
| *Pseudomonas* sp. GV091 | GV091 | GCA_003053805 | A |
| *Pseudomonas* sp. Irchel s3a12 | Irchels3a12 | GCA_900187485 | A |
| *Pseudomonas* sp. Irchel s3f10 | Irchels3f10 | GCA_900187515 | A |
| *Pseudomonas* sp. Irchel s3h9 | Irchels3h9 | GCA_900187475 | A |
| *Pseudomonas* sp. MPR-E5 | MPRE5 | GCA_002883015 | A |
| *Pseudomonas* sp. MS586 | MS586 | GCA_001594225 | A |
| *Pseudomonas* sp. Ok266 | ok266 | GCA_900110195 | A |
| *Pseudomonas* sp. OV221 | OV221 | GCA_003391605 | A |
| *Pseudomonas* sp. OV657 | OV657 | GCA_003386585 | A |
| *Pseudomonas* sp. R15(2017) | R152017 | GCA_002112685 | A |
| *Pseudomonas* sp. R23(2017) | R232017 | GCA_002112575 | A |
| *Pseudomonas* sp. R24(2017) | R242017 | GCA_002113465 | A |
| *Pseudomonas* sp. R27(2017) | R272017 | GCA_002113445 | A |
| *Pseudomonas* sp. Root329 | Root329 | GCA_001424925 | A |
| *Pseudomonas fluorescens* BS2 | BS2 | GCA_000308175 | B |
| *Pseudomonas fluorescens* AHK-1 | AHK1 | GCA_003363095 | B |
| *Pseudomonas fluorescens* FW300-N1B4 | FW300N1B4 | GCA_001625455 | B |
| *Pseudomonas hunanensis* P11 | P11 | GCA_002910975 | B |
| *Pseudomonas marginalis* BS2952 | BS2952 | GCA_900105325 | B |
| *Pseudomonas monteilii* SB3078 | SB3078 | GCA_000510285 | B |
| *Pseudomonas monteilii* SB3101 | SB3101 | GCA_000510325 | B |
| *Pseudomonas monteilii* MO2 | MO2 | GCA_001571445 | B |
| *Pseudomonas plecoglossicida* DJ-1 | DJ1 | GCA_002307455 | B |
| *Pseudomonas plecoglossicida* MR134 | MR134 | GCA_002864885 | B |
| *Pseudomonas plecoglossicida* MR135 | MR135 | GCA_002864795 | B |
| *Pseudomonas plecoglossicida* MR170 | MR170 | GCA_002864845 | B |
| *Pseudomonas plecoglossicida* MR69 | MR69 | GCA_002864775 | B |
| *Pseudomonas plecoglossicida* MR70 | MR70 | GCA_002864905 | B |
| *Pseudomonas plecoglossicida* MR83 | MR83 | GCA_002864865 | B |
| *Pseudomonas plecoglossicida* NyZ12 | NyZ12 | GCA_000831585 | B |
| *Pseudomonas plecoglossicida* TND35 | TND35 | GCA_000764405 | B |
| *Pseudomonas putida* B6-2 | B62 | GCA_000226035 | B |
| *Pseudomonas putida* BIRD-1 | BIRD1 | GCA_000183645 | B |
| *Pseudomonas putida* F1 | F1 | GCA_000016865 | B |
| *Pseudomonas putida* HB3267 | HB3267 | GCA_000325725 | B |
| *Pseudomonas putida* JB | JB | GCA_001767335 | B |
| *Pseudomonas putida* KT2440 | KT2440 | GCA_000007565 | B |
| *Pseudomonas putida* LS46 | LS46 | GCA_000294445 | B |
| *Pseudomonas putida* ND6 | ND6 | GCA_000264665 | B |
| *Pseudomonas putida* S12 | S12 | GCA_000495455 | B |
| *Pseudomonas putida* SJTE-1 | SJTE1 | GCA_000271965 | B |
| *Pseudomonas putida* Idaho | Idaho | GCA_000226475 | B |
| *Pseudomonas putida* CA-3 | CA3 | GCA_002810225 | B |
| *Pseudomonas putida* DPA1 | DPA1 | GCA_002891885 | B |
| *Pseudomonas putida* strain DZ-C18 | DZC18 | GCA_002094795 | B |
| *Pseudomonas putida* FDAARGOS_409 | FDAARGOS409 | GCA_002554535 | B |
| *Pseudomonas putida* H | H | GCA_001077495 | B |
| *Pseudomonas putida* HB13667 | HB13667 | GCA_001306495 | B |
| *Pseudomonas putida* INSali382 | INSali382 | GCA_001653615 | B |
| *Pseudomonas putida* JLR11 | JLR11 | GCA_001183585 | B |
| *Pseudomonas putida* KCJK7911 | KCJK7911 | GCA_003053335 | B |
| *Pseudomonas putida* N1R | N1R | GCA_900156185 | B |
| *Pseudomonas putida* P1 | P1 | GCA_001865225 | B |
| *Pseudomonas putida* PD1 | PD1 | GCA_000799625 | B |
| *Pseudomonas putida* SF1 | SF1 | GCA_001027965 | B |
| *Pseudomonas putida* UV4 | UV4 | GCA_002165695 | B |
| *Pseudomonas putida* UV4/95 | UV495 | GCA_002165665 | B |
| *Pseudomonas putida* TRO1 | TRO1 | GCA_000367825 | B |
| *Pseudomonas taiwanensis* SJ9 | SJ9 | GCA_000500605 | B |
| *Pseudomonas* sp. 22 E 5 | 22E5 | GCA_900004705 | B |
| *Pseudomonas* sp. 2822-17 | 282217 | GCA_002742485 | B |
| *Pseudomonas* sp. 2995-1 | 29951 | GCA_002742505 | B |
| *Pseudomonas* sp. 31 E 5 | 31E5 | GCA_900005815 | B |
| *Pseudomonas* sp. 31 E 6 | 31E6 | GCA_900005935 | B |
| *Pseudomonas* sp. B12(2017) | B122017 | GCA_002113785 | B |
| *Pseudomonas* sp. B13(2017) | B132017 | GCA_002113245 | B |
| *Pseudomonas* sp. B14(2017) | B142017 | GCA_002113255 | B |
| *Pseudomonas* sp. B22(2017) | B222017 | GCA_002113105 | B |
| *Pseudomonas* sp. B23(2017) | B232017 | GCA_002113725 | B |
| *Pseudomonas* sp. B24(2017) | B242017 | GCA_002113085 | B |
| *Pseudomonas* sp. B28(2017) | B282017 | GCA_002113025 | B |
| *Pseudomonas* sp. B4(2017) | B42017 | GCA_002113565 | B |
| *Pseudomonas* sp. B8(2017) | B82017 | GCA_002113575 | B |
| *Pseudomonas* sp. C5pp | C5pp | GCA_000814065 | B |
| *Pseudomonas* sp. CC6-YY-74 | CC6YY74 | GCA_002025205 | B |
| *Pseudomonas* sp. FFUP_PS_41 | FFUPPS41 | GCA_002858645 | B |
| *Pseudomonas* sp. FW305-E2 | FW305E2 | GCA_002901725 | B |
| *Pseudomonas* sp. GTC 16473 | GTC16473 | GCA_001753855 | B |
| *Pseudomonas* sp. GTC 16482 | GTC16482 | GCA_001319995 | B |
| *Pseudomonas* sp. Irchel 3H9 | Irchel3H9 | GCA_900187495 | B |
| *Pseudomonas* sp. JY-Q | JYQ | GCA_001655295 | B |
| *Pseudomonas* sp. Leaf58 | Leaf58 | GCA_001422615 | B |
| *Pseudomonas* sp. LG1E9 | LG1E9 | GCA_003290225 | B |
| *Pseudomonas* sp. MIACH | MIACH | GCA_001269925 | B |
| *Pseudomonas* sp. MR 02 | MR02 | GCA_002797475 | B |
| *Pseudomonas* sp. NBRC 111118 | NBRC111118 | GCA_001320085 | B |
| *Pseudomonas* sp. NBRC 111121 | NBRC111121 | GCA_001320165 | B |
| *Pseudomonas* sp. NBRC 111125 | NBRC111125 | GCA_001320295 | B |
| *Pseudomonas* sp. NBRC 111136 | NBRC111136 | GCA_001320745 | B |
| *Pseudomonas* sp. NBRC 111139 | NBRC111139 | GCA_001753955 | B |
| *Pseudomonas* sp. OV577 | OV577 | GCA_003386595 | B |
| *Pseudomonas* sp. P21 | p21 | GCA_001642705 | B |
| *Pseudomonas* sp. PGPPP2 | PGPPP2 | GCA_002255825 | B |
| *Pseudomonas* sp. RIT357 | RIT357 | GCA_000632245 | B |
| *Pseudomonas* sp. RW405 | RW405 | GCA_003184135 | B |
| *Pseudomonas* sp. SID14000 | SID14000 | GCA_002165135 | B |
| *Pseudomonas* sp. SMT-1 | SMT1 | GCA_003204195 | B |
| *Pseudomonas* sp. SWI36 | SWI36 | GCA_002948105 | B |
| *Pseudomonas* sp. XWY-1 | XWY1 | GCA_002953115 | B |
| *Streptomyces albulus* CCRC 11814 | CCRC11814 | GCA_000403765 | C |
| *Streptomyces albulus* NK660 | NK660 | GCA_000695235 | C |
| *Streptomyces albulus* PD-1 | SaPD1 | GCA_000504065 | C |
| *Streptomyces albus* ZpM | ZpM | GCA_000963515 | C |
| *Streptomyces diastatochromogenes* NRRL B-1698 | NRRLB1698 | GCA_001418405 | C |
| *Streptomyces noursei* ATCC 11455 | ATCC11455 | GCA_001704275 | C |
| *Streptomyces yunnanensis* CGMCC 4.3555 | CGMCC43555 | GCA_900142595 | C |
| *Streptomyces* sp. MspMP-M5 | MspMPM5 | GCA_000373585 | C |
| *Streptomyces* sp. NRRL F-4489 | NRRLF4489 | GCA_001509485 | C |
| *Xenorhabdus bovienii* CS03 | CS031 | GCA_000973125 | D |
| *Xenorhabdus bovienii* CS03 | CS032 | GCA_000973125 | D |
| *Xenorhabdus bovienii* feltiae Florida | Florida | GCA_000736675 | D |
| *Xenorhabdus bovienii* feltiae France | France | GCA_000736655 | D |
| *Xenorhabdus bovienii* feltiae Moldova | Moldova | GCA_000736595 | D |
| *Xenorhabdus bovienii* intermedium | intermedium | GCA_000736615 | D |
| *Xenorhabdus bovienii* kraussei | kraussei | GCA_00073669 | D |
| *Xenorhabdus bovienii* kraussei Quebec | Quebec | GCA_000736555 | D |
| *Xenorhabdus bovienii* oregonense | oregonense | GCA_000736535 | D |
| *Xenorhabdus bovienii* puntauvense | puntauvense | GCA_000736635 | D |
| *Xenorhabdus khoisanae* MCB sp. ICMP 17674 | ICMP17674 | GCA_001037465 | D |
| *Xenorhabdus poinarii* G6 | G6 | GCA_000968175 | D |
| *Xenorhabdus* sp. NBAII XenSa04 | XenSa04 | GCA_000798625 | D |
| Uncultured bacterium | pEAF66 | EU099626 |  |

## REFERENCES

1. Berg G, Grube M, Schloter M, Smalla K. 2014. Unraveling the plant microbiome: looking back and future perspectives. Front Microbiol 5:148.

2. Bais HP, Weir TL, Perry LG, Gilroy S, Vivanco JM. 2006. The role of root exudates in rhizosphere interactions with plants and other organisms. Annu Rev Plant Biol 57:233–266.

3. Arshad M, Frankenberger WT Jr. 1998. Plant growth-regulating substances in the rhizosphere: microbial production and functions. Adv Agron 62:45–151.

4. Handelsman J, Stabb EV. 1996. Biocontrol of soilborne plant pathogens. Plant Cell 8 :1855–1869.

5. Bulgarelli D, Schlaeppi K, Spaepen S, Ver Loren van Themaat E, Schulze-Lefert P. 2013. Structure and functions of the bacterial microbiota of plants. Annu Rev Plant Biol 64:807–838.

6. Lozano GL, Bravo JI, Garavito Diago MF, Park HB, Hurley A, Peterson SB, Stabb EV, Crawford JM, Broderick NA, Handelsman J. 2018. Introducing THOR, a model microbiome for genetic dissection of community behavior. bioRxiv 499715.

7. Valentini M, García-Mauriño SM, Pérez-Martínez I, Santero E, Canosa I, Lapouge K. 2014. Hierarchical management of carbon sources is regulated similarly by the CbrA/B systems in *Pseudomonas aeruginosa* and *Pseudomonas putida*. Microbiology 160:2243–2252.

8. Nishijyo T, Haas D, Itoh Y. 2001. The CbrA-CbrB two-component regulatory system controls the utilization of multiple carbon and nitrogen sources in *Pseudomonas aeruginosa*. Mol Microbiol 40:917–931.

9. Vetter J. 2004. Poison hemlock (*Conium maculatum* L.). Food Chem Toxicol 42:1373–1382.

10. Leete E. 1964. Biosynthesis of the hemlock alkaloids. The incorporation of acetate-1-C_14_ into coniine and conhydrine. J Am Chem Soc 86:2509–2513.

11. Hotti H, Seppänen-Laakso T, Arvas M, Teeri TH, Rischer H. 2015. Polyketide synthases from poison hemlock *(Conium maculatum* L.). FEBS J 282:4141–4156.

12. Roberts MF. 1971. The formation of γ-coniceine from 5-ketooctanal by a transaminase of *Conium maculatum*. Phytochemistry 10:3057–3060.

13. Roberts MF. 1981. Enzymic synthesis of γ-coniceine in *Conium maculatum* chloroplasts and mitochondria. Plant Cell Rep 1:10–13.

14. Schirmer A, Rude MA, Li X, Popova E, del Cardayre SB. 2010. Microbial biosynthesis of alkanes. Science 329:559–562.

15. Blitzke T, Porzel A, Masaoud M, Schmidt J. 2000. A chlorinated amide and piperidine alkaloids from *Aloe sabaea*. Phytochemistry 55:979–982.

16. Abe I, Morita H. 2010. Structure and function of the chalcone synthase superfamily of plant type III polyketide synthases. Nat Prod Rep 27:809–838.

17. Chung EJ, Lim HK, Kim J-C, Choi GJ, Park EJ, Lee MH, Chung YR, Lee S-W. 2008. Forest soil metagenome gene cluster involved in antifungal activity expression in *Escherichia coli*. Appl Environ Microbiol 74:723–730.

18. Davis E, Sloan T, Aurelius K, Barbour A, Bodey E, Clark B, Dennis C, Drown R, Fleming M, Humbert A, Glasgo E, Kerns T, Lingro K, McMillin M, Meyer A, Pope B, Stalevicz A, Steffen B, Steindl A, Williams C, Wimberley C, Zenas R, Butela K, Wildschutte H. 2017. Antibiotic discovery throughout the Small World Initiative: a molecular strategy to identify biosynthetic gene clusters involved in antagonistic activity. Microbiologyopen 6:e00435.

19. Garrido-Sanz D, Meier-Kolthoff JP, Göker M, Martín M, Rivilla R, Redondo-Nieto M. 2016. Genomic and genetic diversity within the *Pseudomonas fluorescens* complex. PLoS ONE 11:e0150183.

20. Thomashow LS, Weller DM. 1988. Role of a phenazine antibiotic from *Pseudomonas fluorescens* in biological control of *Gaeumannomyces graminis* var. tritici. J Bacteriol 170:3499–3508.

21. Haney CH, Samuel BS, Bush J, Ausubel FM. 2015. Associations with rhizosphere bacteria can confer an adaptive advantage to plants. Nat Plants 1:15051.

22. Martins dos Santos VAP, Timmis KN, Tümmler B, Weinel C. 2004. Genomic features of *Pseudomonas putida* strain KT2440, pp. 77–112. In Pseudomonas. Springer US, Boston, MA.

23. Bis DM, Ban YH, James ED, Alqahtani N, Viswanathan R, Lane AL. 2015. Characterization of the nocardiopsin biosynthetic gene cluster reveals similarities to and differences from the rapamycin and FK-506 pathways. ChemBioChem 16:990–997.

24. Peng H, Wei E, Wang J, Zhang Y, Cheng L, Ma H, Deng Z, Qu X. 2016. Deciphering piperidine formation in polyketide-derived indolizidines reveals a thioester reduction, transamination, and unusual imine reduction process. ACS Chem Biol 11:3278–3283.

25. Reynolds T. 2005. Hemlock alkaloids from Socrates to poison aloes. Phytochemistry 66:1399–1406.

26. Hoagland DR, Arnon DI. 1950. The water-culture method for growing plants without soil. Calif Agric Exp Stat Circ 347:1–32.

27. Sivakumar R, Ranjani J, Vishnu US, Jayashree S, Lozano GL, Miles J, Broderick N, Guan C, Gunasekaran P, Handelsman J, Rajendhran J. 2018. Evaluation of InSeq to identify genes essential for *Pseudomonas aeruginosa* PGPR2 corn root colonization. bioRxiv 377168.

28. Hoye TR, Jeffrey CS, Shao F. 2007. Mosher ester analysis for the determination of absolute configuration of stereogenic (chiral) carbinol carbons. Nat Protoc 2:2451–2458.

29. Altschul SF, Gish W, Miller W, Myers EW, Lipman DJ. 1990. Basic local alignment search tool. J Mol Biol 215:403–410.

30. Katoh K, Standley DM. 2013. MAFFT multiple sequence alignment software version 7: improvements in performance and usability. Mol Biol Evol 30:772–780.

31. Sela I, Ashkenazy H, Katoh K, Pupko T. 2015. GUIDANCE2: accurate detection of unreliable alignment regions accounting for the uncertainty of multiple parameters. Nucleic Acids Res 43:W7–14.

32. Kumar S, Stecher G, Li M, Knyaz C, Tamura K. 2018. MEGA X: Molecular evolutionary genetics analysis across computing platforms. Mol Biol Evol 35:1547–1549.

33. Ahmed SA, Lo C-C, Li P-E, Davenport KW, Chain PSG. 2015. From raw reads to trees: Whole genome SNP phylogenetics across the tree of life. bioRxiv 032250.

34. Letunic I, Bork P. 2016. Interactive tree of life (iTOL) v3: an online tool for the display and annotation of phylogenetic and other trees. Nucleic Acids Res 44:W242–5.

